# Predictive models of the genetic bases underlying budding yeast fitness in multiple environments

**DOI:** 10.1101/2025.10.20.683436

**Authors:** Kenia E. Segura Abá, Paulo Izquierdo, Gustavo de Los Campos, Melissa D. Lehti-Shiu, Shin-Han Shiu

**Affiliations:** Genetics and Genome Sciences Graduate Program, Michigan State University, East Lansing, Michigan, United States of America; Department of Energy Great Lakes Bioenergy Research Center, Michigan State University, East Lansing, Michigan, United States of America; Department of Plant Biology, Michigan State University, East Lansing, Michigan, United States of America; Department of Epidemiology and Biostatistics, Michigan State University, East Lansing, Michigan, United States of America; Department of Statistics and Probability, Michigan State University, East Lansing, Michigan, United States of America; Institute for Quantitative Health Science and Engineering, Michigan State University, East Lansing, Michigan, United States of America; Department of Computational Mathematics, Science, and Engineering, Michigan State University, East Lansing, Michigan, United States of America

## Abstract

The ability of organisms to adapt and survive depends on the effects of genes and the environment on fitness. However, the multigenic nature of fitness traits and genotype-by-environment interactions hinder our ability to understand the genetic basis of fitness. Here, we established fitness prediction models for 35 environments using machine learning and existing fitness data and different types of genetic variants for a population of *Saccharomyces cerevisiae* isolates. Models revealed that the predictive ability of genetic variants varied across environments, with copy number variants explaining the majority of fitness variation in most cases. Model interpretation further showed that different variant types identified distinct sets of genes associated with predictive variants. These gene sets were significantly enriched in experimentally validated genes affecting fitness in only a subset of environments, indicating that many genes influencing fitness remain unexplored. Notably, non-experimentally validated genes were more important than validated ones for fitness predictions. Gene contributions to fitness predictions were both isolate and environment dependent, pointing to gene-by-gene and gene-by-environment interactions. Further interpretation of models uncovered experimentally validated and novel candidate genetic interactions for a well characterized stress, the fungicide benomyl. These findings highlight the feasibility of identifying the genetic basis of fitness by using different types of genetic variants and offer novel targets for future functional analysis.

**Author Summary:** Organisms adapt to changing environments by acquiring beneficial traits, which are largely determined by genetic variation. However, predicting how genetic variation influences adaptation, and thus survival, remains a challenge. Here, we used machine learning to identify genes, gene-gene interactions, and gene-by-environment interactions underlying fitness. Specifically, we used machine learning to predict how different genetic variants—such as changes in single nucleotides, presence/absence of a sequence, and differences in copy number—affect fitness in yeast across 35 different environmental conditions. Our results show that prediction accuracy and our ability to interpret the underlying biology depend on the genetic variant type. For example, the best predictions were obtained using differences in copy number. We also found that the contributions of genetic variants to yeast fitness depend on the genetic background. Importantly, our models uncovered known and novel genes that were important across multiple and specific environments and revealed genetic interactions for a well characterized stress, offering insights into how organisms cope with environmental stress. These findings advance our understanding of the genetic basis of fitness and provide a framework for future functional studies and the design of stress-resilient yeast strains.

## Introduction

Deciphering the connection between phenotypic and genetic variation is a long-standing challenge in biology (1). The genetic architecture of phenotypes varies between traits, environments, and populations. Some traits are controlled by a single gene (2) while others are multigenic (3), and most traits are influenced by both genetic and environmental factors (4,5). Furthermore, genetic interactions (e.g., epistasis) and genotype-by-environment interactions also contribute substantially to trait variation (6–15). This complexity frequently results in a non-linear relationship between genotype and phenotype. Thus, prediction of complex traits from genomic data and identification of causal genes remain challenging tasks (16–19).

Quantitative trait locus (QTL) analyses and genome-wide association studies (GWAS) are widely used to uncover genetic variants underlying phenotypic variation (20–23). More recently developed genomic prediction methods (24,25), building upon the principles of QTL analyses, are able to predict complex traits and have led to significant advancements in the fields of crop and animal breeding and human genetics (26,27,16). Examples of successful applications of genomic prediction include accelerating breeding programs for dairy cattle (28,29) and wheat (30) and predicting coronary heart disease risk in humans (31). In contrast to QTL analyses and GWAS, genomic prediction methods based on machine learning are able to capture non-linear relationships between genetic variants and multi-omics data (19,32). For example, transcriptomic data, single nucleotide polymorphism (SNP) genotypes, and methylation data have been used to predict flowering time and to identify interactions between data types (33). Transcriptomic data and SNP genotypes have also been combined with environmental data to predict grain yield in wheat (34) and maize (35). In addition, machine learning-based genomic prediction models can be further interpreted to provide hypotheses of the genetic bases of complex traits (36–39). Such interpretation falls into two main categories: global and local (36). Global interpretation strategies measure the overall contribution of individual predictor variables (features, e.g., a SNP) to trait prediction, whereas local interpretation strategies quantify feature contribution to predicted trait values for each individual in a population. Model interpretation strategies have been used to, for example, understand long non-coding RNA functions in humans (40), identify plant flowering time genes (33), and investigate vertebrate enhancer activity (41).

Interpretable machine learning-based genomic prediction models allow a better understanding of the genotype-to-phenotype association, which has been facilitated by recent population genome sequencing projects. These projects have generated genotype and phenotype data from a relatively larger number of genetically distinct individuals compared to earlier studies in, e.g., human (42–46), *Drosophila melanogaster* (47), *Arabidopsis thaliana* (48–50), and *Saccharomyces cerevisiae* (51–54). In *S. cerevisiae* (budding yeast), a pangenome of 1,011 natural and laboratory isolates is available (51). Furthermore, these isolates have been phenotyped for fitness, a complex trait relevant to adaptation and biotechnology applications, under 35 different environmental conditions (51). They have also been genotyped at single nucleotide polymorphisms (SNPs) and structural variants—e.g., copy number variants (CNVs) and presence/absence variants (PAVs)—have been identified. These types of genetic variants have been shown to be associated with phenotypic variation in various species, including humans, livestock, plants, and yeast (55–62). Together with the rich functional annotation available for budding yeast, the fitness and variant datasets provide a valuable resource for further assessing the impact of environment and types of genetic variants on trait predictions.

Here, we aim to understand the genetic basis of fitness in a natural population of *S. cerevisiae* grown in 35 environments by establishing genomic prediction models using a published dataset (51). Using a subset of 750 diploid isolates, we assessed how well fitness is predicted using SNPs, PAVs, and CNVs in different environments. By interpreting the genomic prediction models, we identified SNP, PAV, and CNV features contributing the most to model performance and quantified their contributions both locally (at the level of individual isolates) and globally in each environment. In addition, we assessed how well our models identified benchmark genes validated in published studies. Finally, we asked what genetic interactions can be discovered by different genetic variant types and how much these interactions contribute to predictions in specific environments.

## Results & Discussion

### Genetic variants differ in their ability to explain variation in fitness across environments

The effects of genetic variants on fitness often depend on the environment, leading to variability in fitness for a single genetic background across environments (51,63). To better understand how fitness responses correlate across environments, we examined fitness values (colony size relative to that in a control environment) of a population of 750 diploid budding yeast isolates in 35 environments (51). An environment is defined as a treatment with a specific temperature or a chemical at a specific concentration (**S1 Table**). Pearson’s correlation coefficients (*r*) were estimated between fitness values of pairs of environments and clustered using Euclidean distance (**Fig 1A**). We found eight environment clusters (*r* > 0.51) encompassing 24 of 35 environments (**Fig 1A**). Cluster 1 was indicative of similar cellular response mechanisms—anisomycin and cycloheximide inhibit peptide chain elongation by binding to different subunits of the ribosome (64,65). Environments within the remaining clusters except cluster 8 tended to be similar temperatures (cluster 2: 40°C and 42°C), to be the same chemicals but at different concentrations (clusters 3, 4, and 7: caffeine, benomyl, and formamide, respectively), or to have shared chemical properties (cluster 5: xylose, ribose, sorbitol, glycerol, and ethanol; cluster 6: LiCl and NaCl). Although the environments in cluster 8 do not have obvious relationships, there are likely similar genetic mechanisms underlying differences in fitness among isolates in these environments.

**Fig. 1.**
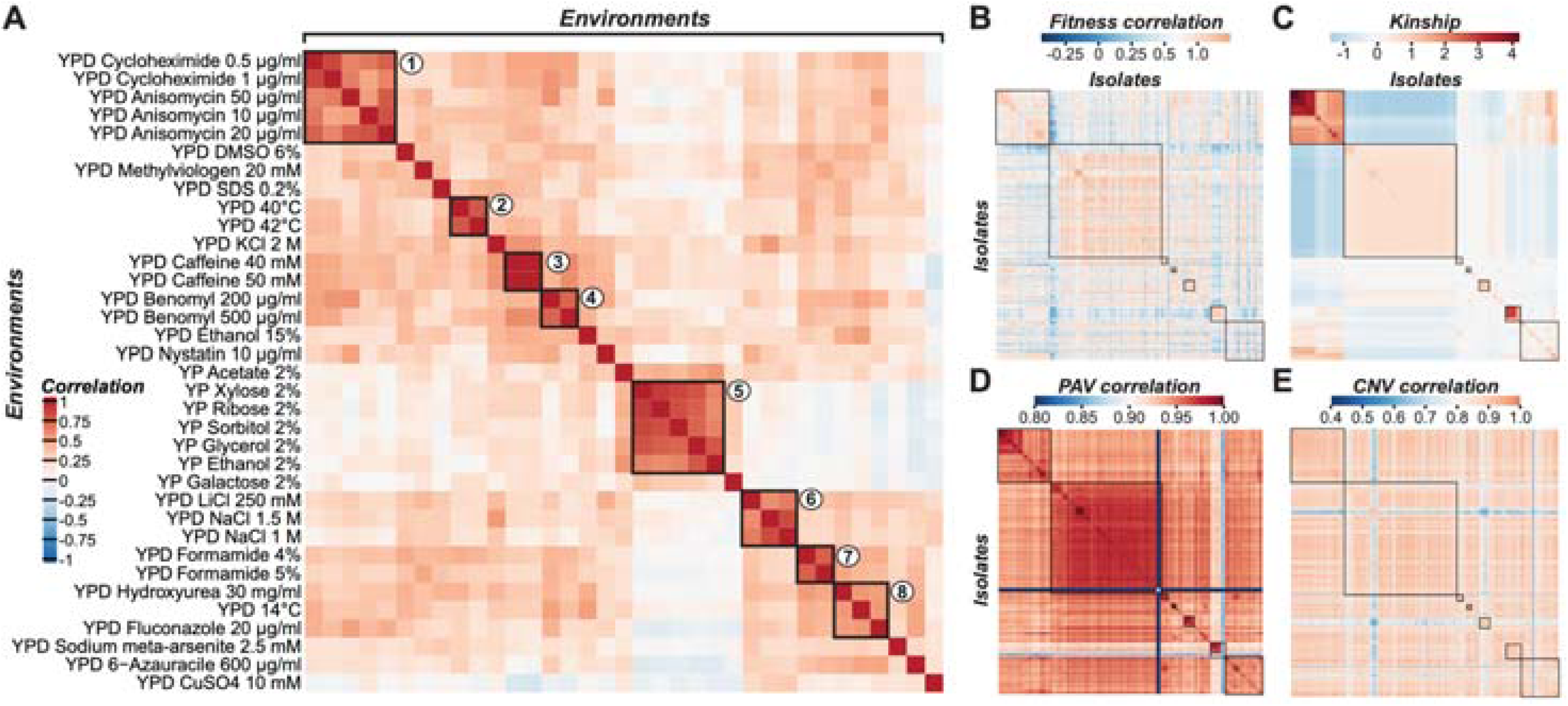
Phenotypic correlation between environments based on fitness values and relationships between yeast isolates based on fitness and genotype data. **(A)** Relationships between fitness values for pairs of environments shown as Pearson’s correlation coefficients (correlation). Colors indicate degrees of correlation. Clusters (rectangles labeled 1–8) were determined via hierarchical clustering based on the Euclidean distances of yeast isolate fitness values across environments. **(B–E)** Relationships between 750 diploid yeast isolates based on **(B)** correlations of fitness across 35 environments, **(C)** kinship (calculated using SNPs), **(D)** correlations of PAV profiles, and **(E)** correlations of CNV profiles. Kinship was used to order isolates and identify clusters of isolates (black boxes).

While the presence of environment clusters is expected because isolates respond similarly to related treatments, there are substantial differences in fitness values between isolates across environments (**Fig 1B**), raising the question of the extent to which genetic variation across budding yeast isolates explains fitness differences across environments. To address this, we calculated the correlation between relatedness estimated using three types of genetic variants—SNPs (i.e., kinship, **Fig 1C**), PAVs (**Fig 1D**), and CNVs (**Fig 1E**)—and fitness profiles of isolates across environments. The kinship matrix reveals clusters indicative of population structure (rectangles, **Fig 1C**) that are partially preserved in a matrix of fitness correlations (**Fig 1B**), indicating that the variation in fitness across environments is correlated with population structure. This pattern also holds for PAV profile correlations (**Fig 1D**), but it is not as clear for CNV profiles (**Fig 1E**). Thus, fitness is expected to be more highly correlated with kinship (Spearman’s rank ρ = 0.27, *p* < 2.2×10^-16^, **S1A Fig**) and PAV correlations (ρ = 0.30, *p* < 2.2×10^-16^, **S1B Fig**) than with CNV correlations (ρ = 0.14, *p* < 2.2×10^-16^, **S1C Fig**). Furthermore, kinship was highly correlated with PAV correlations (ρ = 0.48, *p* < 2.2×10^-16^, **S1D Fig**) but not with CNV correlations (ρ = 0.10, *p* < 2.2×10^-16^, **S1E Fig**), and PAV correlations and CNV correlations were lowly correlated (ρ = 0.17, *p* < 2.2×10^-16^, **S1F Fig**). These findings indicate that different genetic variant types may contain overlapping and distinct information that can be used to predict fitness in different environments.

### Genetic variant type influences model performance in fitness prediction

To determine how well different types of genomic variants predict yeast fitness in a given environment, we built single-environment fitness prediction regression models using five algorithms with SNPs, PAVs, or CNVs as input features (see **Materials and Methods**). For each environment and algorithm, “baseline” models were constructed using the first five principal components (PCs) of the SNP data as a proxy of population structure (capturing 59% of the genetic variation), and “optimized” models were generated using a feature selection approach to maximize prediction accuracy (see **Materials and Methods**). Model performance was assessed by calculating the coefficient of determination between true and predicted fitness values (R^2^, **S2 Table**). Random Forest (RF) testing set performances tended to be better than those of the other algorithms (**S3 Table**).

Different types of genetic variants have distinct, environment-dependent contributions to fitness prediction. Regardless of algorithm, the R^2^ of models based on population structure alone varied greatly between environments, ranging from 0 to 0.56 (PCs, median R^2^ = 0.15 and mean R^2^ = 0.20, **Fig 2A**, **S2 Table**). In 32 environments, SNP, PAV, and/or CNV information improved models compared with those based solely on population structure (**Fig 2A**).

**Fig 2.**
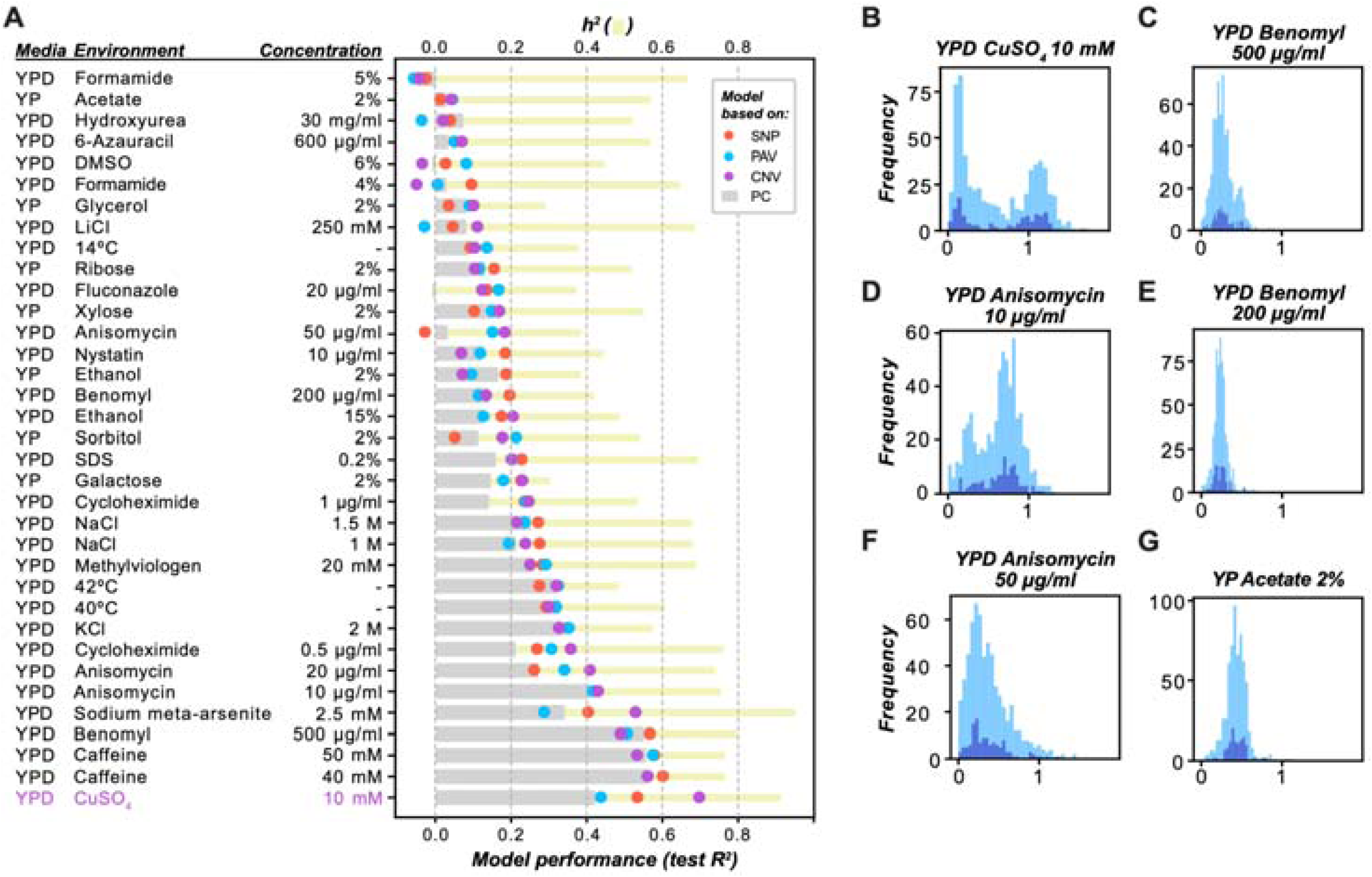
Narrow-sense heritability, model performance, and fitness distributions across environments. **(A)** Narrow-sense heritability (*h^2^*; yellow bars) estimates and the test set performances (R^2^) of the optimized Random Forest models for each environment. Single-environment models were built with PCs (approximate population structure; gray bars), SNPs (orange), PAVs (blue), or CNVs (purple). The YPD CuSO_4_ 10 mM model using CNV features achieved the highest test performance (purple). **(B–G)** Distribution of fitness values for the **(B)** YPD CuSO_4_ 10 mM, **(C)** YPD Benomyl 500 μg/ml, **(D)** YPD Anisomycin 10 μg/ml, **(E)** YP Benomyl 200 μg/ml, **(F)** YPD Anisomycin 50 μg/ml, and **(G)** YPD Acetate 2% environments. Light and dark blue histograms represent fitness value distributions of the training and test sets, respectively.

Surprisingly, despite the correlation between CNV and fitness variation being lower than that between SNP or PAV and fitness (**Figs S1A**, **S1B**, and **S1C**), the CNV-based model performance (median R^2^ = 0.21) was similar to that using SNPs (median R^2^ = 0.20) or PAVs (median R^2^ = 0.18). This seemingly contradictory result is likely due to the higher degree of dependence of SNPs and PAVs on population structure relative to that of CNVs, suggesting that CNVs may identify candidate fitness genes that differ from those identified by population structure. Utilizing CNVs to identify candidate fitness genes is feasible because they have been reported to have large deleterious effects on fitness that may also vary depending on the genetic background and the environmental condition (51,59,62,66,67). Furthermore, CNVs were found to explain more trait variation than SNPs on average (51), which may also explain the similar or slightly higher performance of CNV-based models compared to the SNP- and PAV-based models in certain environments.

Using models established with the RF algorithm, fitness values in three environments (YPD Caffeine 40 mM, YPD Caffeine 50 mM, and YPD Benomyl 500 μg/ml) were predicted well by three feature types (SNPs, PAVs, and PCs) (**Fig 2A**), and the highest prediction performance was observed for YPD CuSO_4_ 10 mM with CNV features (R^2^ = 0.70, **Fig 2A**). SNP-, PAV-, CNV-, and PC-based models performed the best in 34.2% (12 out of 35), 22.9% (8), 37.1% (13), and 5.7% (2) of the environments, respectively. CNV-based models outperformed SNP-, PAV-, and PC-based models in 51.4% (18), 51.4%, and 62.9% (22) of environments, respectively. In particular, CNV-based models outperformed the PC-based models by 1.66-fold for CuSO_4_ 10 mM and 1.56-fold for sodium meta-arsenite 2.5 mM. The higher performance of CNV-based models in certain environments may reflect the fact that CNVs lead to variation in the dosage of genes important for growth under stress conditions and may provide a selective advantage (62). For example, haploid budding yeast strains with >1 copy of *CUP1* (copperthionein) have higher fitness than those that have only one copy (68). On the other hand, SNP- and PAV-based models outperform CNV-based models in environments such as YPD NaCl 1.5 M and YPD Methylviologen 20 mM (**Fig 2A**), suggesting that SNPs and PAVs likely contribute more to fitness in these environments than CNVs do. Whether this reflects underlying genetic mechanisms remains unclear. Lastly, a large proportion of the trait heritability in most environments remains unexplained by any individual variant type (**Fig 2A**).

To assess why fitness is better predicted in some environments than in others, we established a linear model predicting the performances of the optimized single-environment RF models with three features—narrow-sense heritability (*h^2^*) of fitness, fitness variance, median fitness—along with all pairwise and three-way interaction terms (see **Materials and Methods**). These three features explained 42% (i.e., adjusted R^2^ = 0.42), 45%, 33%, and 59% of the variation in performance for PC, SNP, PAV, and CNV-based models, respectively (**S4 Table**). Different combinations of these features contributed to PC, SNP, PAV, and CNV model performances to varying degrees. PAVs and CNVs were better at predicting fitness traits with higher variance, lower median fitness, and lower fitness variance-by-*h^2^* terms; PCs were predictive of environments with lower fitness variance-by-*h^2^* terms; and no terms were statistically significantly correlated with SNP model performances (for statistics and term ranges, see **S4 Table**). To better understand which feature(s) are important for model performance, we determined SHapley Additive exPlanations (SHAP) values based on the same linear models, where positive values indicate that a feature increased model performance and negative values indicate that a feature decreased performance relative to the expected performance of a model trained on one environment. Based on SHAP values, *h^2^* was the most correlated with model performances out of the fitness-related features tested (PC: *r* = 0.62, SNP: *r* = 0.64, PAV: *r* = 0.54), but it was negatively correlated with CNVs (*r* = -0.67, **S2 Fig**). Since *h^2^* was estimated from SNPs, this result is consistent with our finding that SNP profiles are correlated with PAVs but not with CNVs (**Figs S1D** and **S1E**). Because PCs are also derived from SNPs, and population structure confounds the relationship between PAVs and SNPs (**Figs 1C**, **1D**, and **S1D**), the correlation between *h^2^* and PC-, SNP-, and PAV-based model performances may be driven by shared genetic information.

A factor that may influence model performance is the genetic architecture of the trait, which can be partially assessed based on the shape of the fitness distribution (e.g., number of peaks or skewness). For example, traits with bimodal distributions tend to have Mendelian inheritance (69), which could be easier to predict. Consistent with this, models of environments with the highest performances, such as YPD CuSO_4_ 10 mM, YPD Benomyl 500 μg/ml, and to a lesser extent, YPD Anisomycin 10 μg/ml, exhibited bimodal distributions of fitness (**Fig 2B**, **2C**, and **2D**), whereas other environments with non-bimodal distributions of fitness, such as YPD Benomyl 200 μg/ml (**Fig 2E**), YPD Anisomycin 50 μg/ml (**Fig 2F**), and YP Acetate 2% (**Fig 2G**), were not predicted well. However, YPD Caffeine 40 mM and 50 mM were predicted well despite having non-bimodal distributions (**S3 Fig**). In addition to fitness distributions, the number of features used to train models also influences model performance. Feature number explained 33% (adjusted R^2^, coefficient = 7.2×10^-5^, *P* = 1.7×10^-4^, **S4 Table**), 3%, and 0% of the variation in SNP-, PAV-, and CNV-based model performances, respectively. Taken together, these results indicate that model performance arises from a combination of technical factors, such as feature number, and genetic mechanisms and interactions that influence trait distribution and trait variance, with their contributions differing across genetic variant types.

### Different genetic variant types and environments uncover distinct sets of genes predictive of fitness

To further assess the genetic basis of fitness in an environment, we determined which SNP, PAV, or CNV (i.e., features) variants contribute to fitness predictions in the optimized models for different environments using two feature importance measures—Gini importance (70) and SHAP (39,71). Gini importance provides a global, overall measure of feature importance among all yeast isolates, whereas SHAP values allow further exploration of feature contributions to fitness in each yeast isolate. Gini- and average absolute SHAP value-based feature rankings were significantly correlated with each other in each of the five optimized RF models with the highest performances for any genetic variant type (**S4A Fig**, Spearman’s correlation coefficients for RF models trained on complete feature sets: **S4B Fig**, **S5 Table**). Using the five environments where the optimized models have the best performance (**Fig 2**) as examples, SHAP value-based rankings of shared features between PAVs and CNVs (i.e., open reading frames [ORFs] that were important for predictions in both models for an environment) were significantly correlated for four environments (*p* `≤ 0.001,` **Fig 3A**, Spearman correlations based on Gini importance: **S4C Fig**, Spearman correlations for RF models trained on complete feature sets: **Figs S4D** and **S4E**, **S6 Table**). This overlap is expected since PAV and CNV are structural variants that are partially dependent on each other; however, there remains a substantial number of non-overlapping, predictive ORF features. There was little overlap in the feature importance rankings of the best five environments when comparing SNP vs PAV models and SNP vs CNV models, regardless of the feature importance measure examined (for optimized models: 3.6×10^-2^≤ *p* ≤ `0.9, for complete models: 1.3×10`^-15^ ≤ *p* ≤ `1.0`, **Figs 3A**, **S4C**, **S4D**, and **S4E**, **S6 Table**).

**Fig 3.**
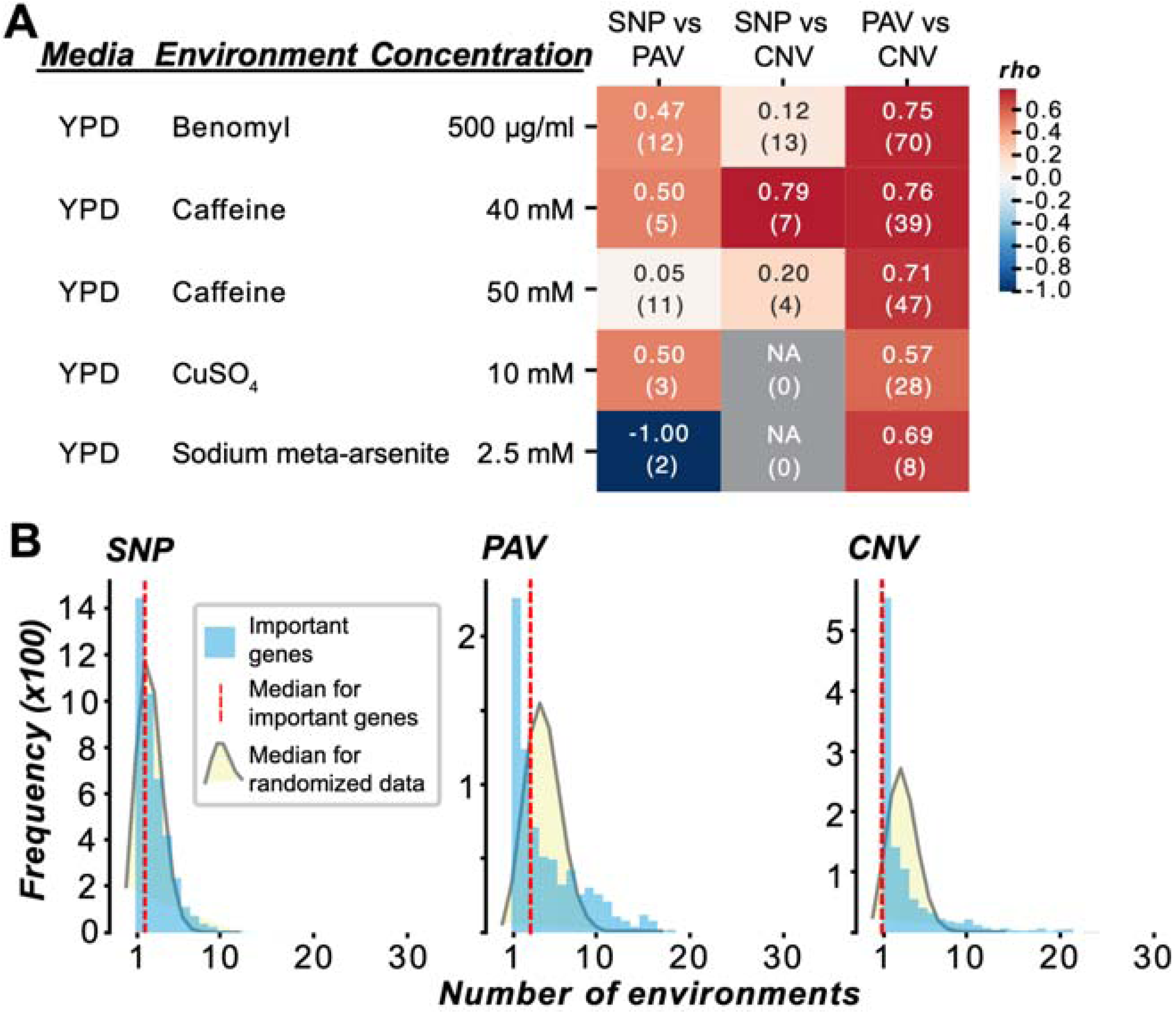
Comparisons of feature importance between variant types and across environments. **(A)** Spearman’s rank correlation (rho) between the average absolute SHAP values from the optimized RF models trained on different genetic variant types (e.g., SNP vs PAV optimized RF models’ SHAP values were compared) for the five best predicted environments. The number of genes that were shared by both feature sets is denoted in parentheses. Gray boxes indicate there was no overlap in important genes between feature sets. **(B)** Average absolute SHAP values from the optimized RF models were used to determine the distribution of unique or shared genes with non-zero importance across 1 to 35 environments (blue bars; red line: median). This distribution was compared to a null distribution of median randomized counts (see **Materials and Methods**) using the Kolmogorov-Smirnov test (alternative = “greater”); for SNPs: median *P* = 6.1×10^-28^; for PAVs: *P* = 1.5×10^-31^; for CNVs: *P* = 2.1×10^-71^.

The lack of overlap between SNPs and CNVs may stem from their different associations with genes. CNVs can span small or large segments of DNA and even entire chromosomes (72). They can alter not only protein-coding but also regulatory regions, leading to changes in gene expression levels, which in turn can influence the regulation of other genes and downstream phenotypes, including fitness (62). On the other hand, SNPs are point mutations that may simply be linked to the causal variants and may have a smaller impact compared to CNVs. Considering the differences in the sets of genes uncovered by these three types of features, future studies on trait variation may benefit from considering multiple types of genetic variants, in addition to SNPs, to obtain a more complete picture of the mechanisms underlying trait variation.

Our finding that related environments cluster together using fitness data (**Fig 1A**) prompted us to ask if similar genetic mechanisms underlie fitness variation in related environments. To test this, we examined the overlap of predictive features across models using SHAP-based feature rankings. Among the five optimized RF models with the highest performances, YPD Caffeine 40 mM and 50 mM, which clustered together based on fitness values (**Fig 1A**), shared 578 SNPs, 64 PAVs, and 55 CNVs (**S7 Table**). The rankings of these features between the two environments were significantly correlated at different strengths depending on the genetic variant (SNP: Spearman’s □ = 0.44, *P* = 4.91×10^-29^; PAV: □ = 0.85, *P* = 3.00×10^-19^; CNV: □ = 0.74, *P* =1.15×10^-10^), indicating that similar genetic mechanisms underlie fitness in these two environments. However, when examining the overall overlap of predictive genes across all 35 environments, we found that significantly fewer predictive genes were shared across environments than expected by random chance (**Fig 3B**, **S4F Fig**), pointing to the impact of genotype-by-environment interactions and the largely unique genetic bases for fitness.

Having identified important, predictive genes in each environment, we next asked what functions these genes have that may be relevant to the environments in which they are important for fitness predictions. To explore this, we conducted gene ontology (GO) and pathway enrichment analyses of the important genes from the optimized RF models (**S8 Table**, see **Materials and Methods**). We found no significantly enriched GO terms in the five optimized, best-performing RF models, but found a total of 10 enriched GO terms in eight of the remaining environments, for models using CNVs or PAVs. At the pathway level, we identified a total of 16 enriched pathways among 11 environments (**S8 Table**). For example, in the Sodium meta-arsenite 2.5 mM model, CNV features were enriched for genes related to arsenate detoxification (odds ratio = 166.7, *q* = 0.01) and also pyridoxal 5’-phosphate biosynthesis II pathways (odds ratio = 249.7, *q* = 0.01), which are necessary for arsenic resistance (73). SNP features for the YPD Caffeine 40 mM model were enriched for glutamine degradation I genes (odds ratio = 6.5, *q* = 0.005) and glutaminyl-tRNA^gln^ biosynthesis via transamidation genes (odds ratio = 6.5, *q* = 0.005). No functional connections between caffeine and glutamine degradation or glutamyl-tRNA biosynthesis were found in the literature. However, the fact that CNVs identified genes related to arsenic resistance highlights the importance of considering structural variants to understand better the genetic basis of fitness in response to arsenic stress, and likely other complex traits.

To further explore the functions of the genes predicted as important for fitness in specific environments, we assessed whether genes experimentally verified to be important for survival in an environment (benchmark fitness genes) were enriched among the important genes in the five best predicted environments—YPD Caffeine 40 mM, YPD Caffeine 50 mM, YPD Benomyl 500 μg/ml, YPD CuSO_4_ 10 mM, and YPD Sodium meta-arsenite 2.5 mM (**Fig 2A**, **S9 Table**). Benchmark fitness genes for benomyl, caffeine, copper(II) sulfate, and sodium meta-arsenite were collected from the *Saccharomyces* Genome Database (SGD) or manually curated from the literature (see **S10 Table**). Manually curated benchmark genes were significantly enriched for YPD Sodium meta-arsenite 2.5 mM using CNVs in the top 1% of genes (**S11 Table**). SGD copper benchmark fitness genes were significantly enriched in the top 5% of SNPs from the YPD Sodium meta-arsenite 2.5 mM model (**S11 Table)**. No benchmark gene enrichment was observed in the top 10% of features for any model (**S11 Table)**. SGD caffeine benchmark genes were significantly enriched in the top 15% and 20% of SNP features from both the YPD CuSO_4_ 10 mM and YPD Caffeine 50 mM models and in the top 25% of SNP features from the YPD Caffeine 40 mM and 50 mM models (**S11 Table**). These benchmark gene enrichment analyses confirm that predicted genes are functionally relevant to fitness in their respective environment (i.e., caffeine benchmark genes enriched in the YPD Caffeine 50 mM model’s important features). However, the general lack of benchmark gene enrichment in important genes (above the 90th percentile) of relevant models prompted us to consider possible explanations.c

### Non-benchmark genes explain more fitness variation than benchmark genes

There are three potential reasons for the limited enrichment of environment-specific and cross-environment benchmark fitness genes. First, some of the important genes contributing to predictions may be false positives. While the R^2^ was as high as ∼0.7, the models are far from perfect, and false predictions are expected. Second, the fitness trait distributions suggest complex genetic bases (**Fig 2B**), and genes with smaller fitness effects are inherently harder to verify experimentally. Thus, it is likely that a substantial number of relevant genes are not present in the benchmark sets. Third, the genetic background in which benchmark fitness genes were experimentally verified was limited to laboratory strains, particularly S288C and W303, the former of which is absent from the diploid budding yeast population dataset. Since laboratory strains have undergone extensive selection under controlled conditions, benchmark genes, which are predominantly discovered in the lab strains, may not all be important for fitness in natural isolates.

If the first possibility is true, we expect that, when a model is trained with features devoid of benchmark gene sets, the model performance should decrease appreciably because the predictive power is mainly derived from benchmarks in the training data, rather than the important, non-benchmark genes we identified. Conversely, if the second possibility is true, i.e., there remains a substantial number of relevant genes not covered by the benchmark genes, then the models devoid of benchmarks would perform just as well as, if not better than, the models including benchmarks. To test this, a new set of RF models were trained using feature sets consisting of only benchmark genes, only important non-benchmark genes, or both benchmark and important non-benchmark genes (see **Materials and Methods**). Models trained on the feature set consisting of both benchmark and non-benchmark genes (“combined models”, **Fig 4A**) or those consisting of only important non-benchmark genes (**Fig 4B**) performed better than models built using benchmark genes alone (**Fig 4C**) when using PAVs and CNVs. Differences in test R^2^ between the important non-benchmark gene models or the combined models and the benchmark gene models ranged from 0.07 to 0.62 and 0.08 to 0.61, respectively. There were less performance differences between the SNP-based models (differences in test R^2^ between important non-benchmark gene models or the combined models and the benchmark gene models ranged from 0 to 0.05 and 0.01 to 0.06, respectively, **Fig 4A-C**). These results indicate that structural variation in non-benchmark genes explains more variation in fitness than benchmark genes across the selected environments. It is possible that the better or similar performances of important non-benchmark gene-based models are due to more features being used, but there was no significant association between the number of features used for training and model performance (**S12 Table**), making this possibility unlikely. Taken together, our results suggest that the limited enrichment of benchmark genes is not likely due to the first possibility (predicted important genes are false positives) and support the second possibility, where the important non-benchmark genes explain a high proportion of variance in fitness.

**Fig 4.**
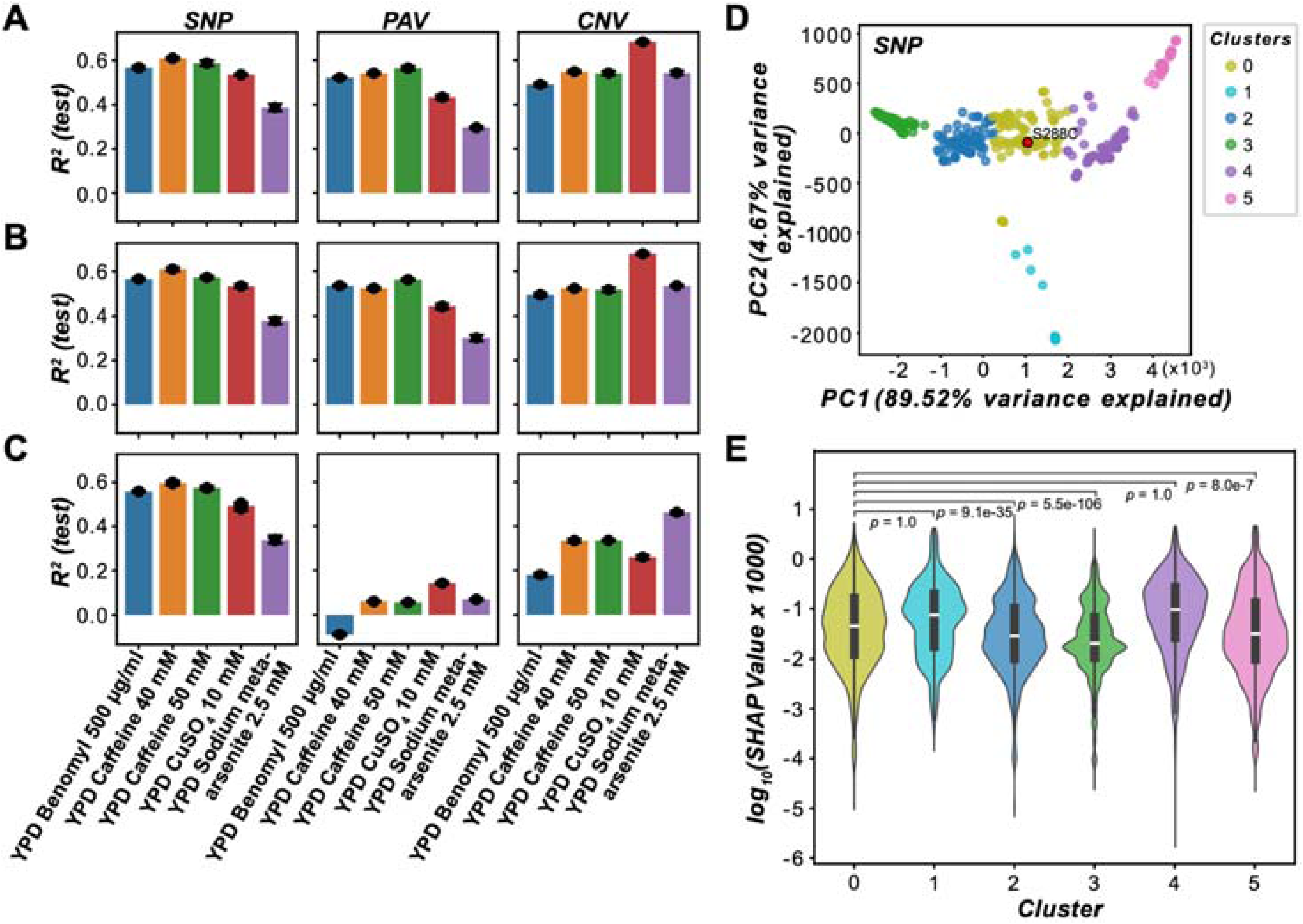
Contribution of benchmark genes to fitness predictions, as well as comparisons of SHAP values between clusters of isolates. **(A–C)** Performance of RF fitness prediction models built with different feature sets to predict fitness in YPD Benomyl 500 μg/ml (blue), YPD Caffeine 40 mM (orange), YPD Caffeine 50 mM (green), YPD CuSO_4_ 10 mM (red), and YPD Sodium meta-arsenite 2.5 mM (purple). Feature sets consisted of **(A)** both important non-benchmark genes identified by the optimized RF models and benchmark genes, **(B)** only important non-benchmark genes, or **(C)** only benchmark genes. **(D)** Principal component analysis was performed on the Euclidean distance matrix calculated from the SNP genotypes to assess genetic relatedness among isolates. Isolates are colored according to memberships in six K-means clusters of similar isolates identified using the same distance matrix. **(E)** Violin plot of the distributions of the absolute SHAP values from the optimized YPD Caffeine 50 mM SNP model for the clusters of isolates identified in (D). *P*-values are from Mann-Whitney U tests conducted to assess the relationship of median absolute SHAP values of benchmark genes between clusters.

Next, we reasoned that if the third possibility is true, i.e., benchmarks predominantly discovered in laboratory strains like S288C are less important for fitness in wild yeast isolates, then the feature importance (absolute values of the SHAP values) of benchmark fitness genes should decrease for yeast isolates that are increasingly genetically distant to laboratory strains. To test this, we first clustered the isolates based on their genetic distance, identifying a cluster containing S288C (yellow cluster 0; SNP: **Fig 4D**; PAV: **S5 Fig**) and other clusters with varying genetic distances to S288C (non-yellow clusters; **Figs 4D** and **S5**), and then compared the SHAP values from the optimized RF models for benchmark genes between clusters. Using the SNP-based YPD Caffeine 50 mM model as an example, the caffeine benchmark gene SHAP values were significantly greater for isolates more related to S288C (cluster 0) than for the more distantly related isolates in clusters 2 (*P* = 9×10^-35^), 3 (*P* = 5×10^-106^), and 5 (*P* = 8×10^-7^), but not clusters 1 and 4 (*P* = 1, **S13 Table**, **Fig 4E**). For the PAV-based models, only two or fewer benchmark gene features were found among the important features; thus, no comparisons of SHAP values were made between clusters. The association using the SNP models provides support for the third hypothesis that the bias in the identification of benchmark fitness genes from specific genetic backgrounds, such as laboratory strains, leads to an underestimate of model performance for certain environments. In addition, many genes that were among the important features from the optimized PAV models for the top five best-predicted environments were absent in S288C (mean = 67.7%, sd = 4.8%), partially explaining the lack of enrichment of benchmark fitness genes predicted by PAV models.

### Fitness effects of genetic variants are isolate-dependent

The feature importance analyses reported in earlier sections (e.g., **Fig 3**) involve interpreting models globally, i.e., the importance of a variant or a gene indicates its average contribution to fitness predictions among isolates. This raises the question whether the genes that contributed the most to the model predictions do so across most if not all isolates or in an isolate-dependent manner. To address this, we interpreted feature contributions locally, i.e., at the level of individual isolates, using SHAP values from the optimized SNP, PAV, and CNV RF models for the environments with the five best predictions. Isolates were clustered based on the SHAP values of the top 20 most predictive features in each model (see **Materials and Methods**). Clustering of SHAP values across isolates revealed that the feature importances significantly correlated with fitness in 14 out of 15 models, indicating isolate-dependent variant contributions to fitness predictions (SNP: five environments, PAV: five, CNV: four, **S14 Table**). For example, SNP-based median absolute SHAP values showed the strongest positive correlation with fitness in YPD Caffeine 40 mM (*slope* = 306.8, *P* = 1×10^-52^, **S14 Table**, **S6 Fig**), followed by YPD Benomyl 500 μg/ml (*slope* = 273.57, *P* = 2×10^-42^, **Figs 5A** and **5B**, **S14 Table**). Similar trends were observed with other variant types, although not as strongly as with SNPs (PAVs: *slope* = -49.2–94.2; CNVs: *slope* = 34.7–87.0, **S14 Table**).

**Fig 5.**
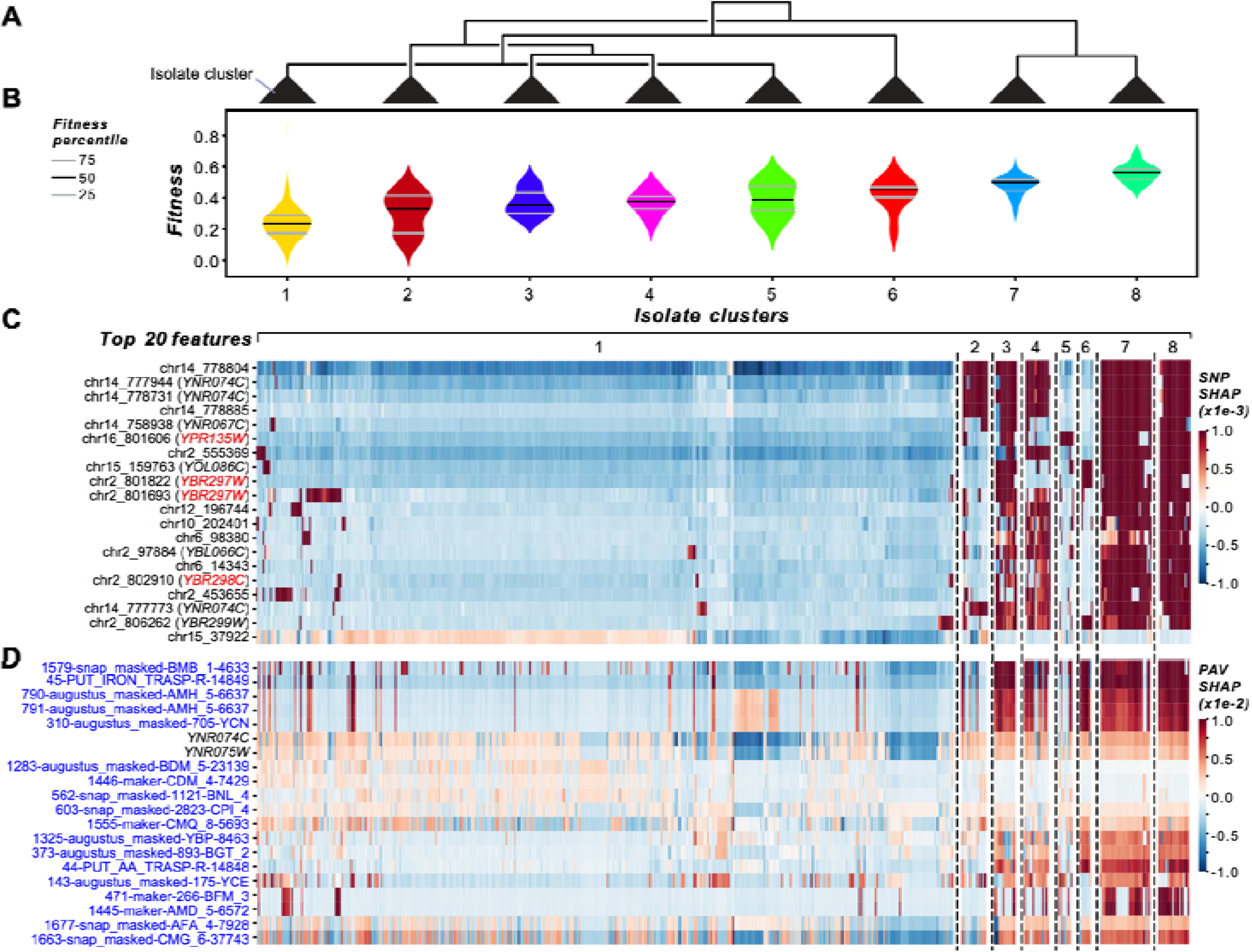
Importance of variants in predicting fitness effects in different isolates. **(A)** Dendrogram showing clusters of isolates based on the SHAP values of the top 20 SNP features from the YPD Benomyl 500 μg/ml optimized model. Dendrogram triangles denote different clusters. **(B)** Violin plot of fitness distributions of isolates in each cluster identified in (A). **(C)** Heatmap of SHAP values of the top 20 SNP features corresponding to the dendrogram in (A). Benchmark benomyl genes are colored in red on the left side of the heatmap. **(D)** Heatmap of SHAP values of the top 20 PAV features (blue text). Black text: PAV features that mapped to genes. Isolates are ordered based on the SNP-based isolate clusters.

Next, we focused on YPD Benomyl 500 μg/ml for interpretation because it included benchmark genes among the top 20 most predictive SNP features (**Fig 5C**). From the eight clusters identified from the SNP-based SHAP values (**Figs 5A**, **5B**, and **5C**), cluster 1 had the lowest median fitness, and most of the top features had negative SHAP values, indicating that these variants contributed to decreased predicted fitness for isolates in cluster 1. In contrast, the same features tended to have positive SHAP values in clusters with higher fitness (clusters 7 and 8). Clusters 2, 3, and 4 showed more heterogeneous SHAP profiles with combinations of both positive and negative contributions that correlated with intermediate to low fitness. Clusters 5 and 6 defy the above generalization but contain a small number of genes with positive SHAP values that are also benchmark benomyl genes (*YPR135W* and *YBR297W*, red feature names, **Fig 5C**). *YPR135W* (*CTF4*, Chromosome transmission fidelity) is required for sister chromatid cohesion (74). Mutations in various *CTF* genes were found to exhibit both tolerant and sensitive growth phenotypes in yeast grown under benomyl stress (75). *YBR297W* (*MAL33*, Maltose activator) encodes the transcriptional activator that regulates the expression of *MAL31* (maltose permease) and *MAL32* (maltase), which are required for maltose fermentation (76). Although no direct connection between *MAL31* and benomyl was found, benomyl has been shown to positively affect desirable traits of aneuploid wine-making yeast strains (77). It would be interesting to investigate the potential effects of benomyl on maltose fermentation of diploid *S. cerevisiae* strains.

The SHAP values of the top 20 PAV (**Fig 5D**) and CNV features (**S7 Fig**) generally followed similar clustering patterns as the SNPs but with more admixture. The top 20 PAV and CNV features tended to contribute to lower predicted fitness in isolates from clusters 1 and 2, while a subset of features showed more positive SHAP values across isolates with higher predicted fitness. In clusters 3, 4, 6, 7, and 8, PAV and CNV features generally had more positive SHAP values, but this pattern was inconsistent with the fitness of the SNP-based isolate clusters, where isolates in clusters 3 and 4 tended to have lower fitness and isolates in clusters 6, 7, and 8 tended to have higher fitness. This finding indicates that the PAV and CNV SHAP values do not fully reflect the fitness patterns associated with SNP-based SHAP clusters, highlighting the importance of analyzing multiple variant types to understand the genetic basis of fitness. Furthermore, there was minimal overlap in the genes among the top 20 SNPs and PAVs or CNVs, indicating that each variant type uncovers distinct aspects of the genetic basis of fitness variation across isolates.

We observed that no single feature was overwhelmingly important for predicting fitness in most environments, underscoring the polygenic basis of fitness in these environments (**Figs 5**, **S6**, **S7**, **S8**, **S9**, and **S10**). The most notable exception was the YPD CuSO_4_ 10 mM CNV model, where fitness variation is driven mainly by *CUP1-2* (*YHR055C*, **S8 Fig**), the major gene controlling copper toxicity response in yeast (76). Another notable exception is an ORF absent in the S288C reference genome (1594-snap_masked-AMH_5-6573) that is a major contributor to higher fitness in the YPD Caffeine 40 mM environment for one cluster of isolates (cluster 8 in **S6 Fig**). The closest BLASTx and tBLASTx match for this ORF (percent identity = 96.93, E-value = 0, **S15 Table**) is the gene *YLR342W* (*FKS1*, FK506 Sensitivity), which encodes the catalytic subunit of 1,3-β-D-glucan synthase, which is found in the cell wall of most fungi (78). It remains unclear how this gene may impact response to caffeine. Lastly, in the PAV-based YPD Sodium meta-arsenite 2.5 mM model, *YGL258W* (*VEL1*, Velum formation), a protein with unknown function in budding yeast, contributed to lower predicted fitness in a subset of isolates, whereas the corresponding CNV model identified several genes known to confer tolerance to arsenate when overexpressed (79), including *ARR1* (Arsenicals resistance, *YPR199C*), *ARR2* (*YPR200C*), *ARR3* (*YPR201W*), *YPR196W* (Putative maltose-responsive transcription factor), and *YPR198W* (*SGE1*, Suppression of Gal11 expression), as contributing to higher predicted fitness in certain isolates (**S9 Fig**). Taken together, our findings indicate that model interpretation based on SHAP values allows identification of genetic variants important for predicting fitness in different isolates and hypothesizing which stress-response genes drive fitness differences among isolates. It is also notable that the majority of features within the top 20 are either intergenic SNPs or ORFs not found within the reference genome (S288C), providing an opportunity for further experimental exploration.

### Genetic interactions underlying fitness variation

We found that multiple important features that are predictive of fitness in an environment show similar SHAP value patterns within the same genetic backgrounds (**Figs 5C**, **5D**, **S6**, **S7**, **S8**, **S9**, and **S10**), indicating that they contribute similarly to fitness predictions, potentially via shared genetic mechanisms or genetic interactions. To assess the contribution of genetic interactions to fitness predictions, we first asked if experimentally validated genetic interactions are predictive of fitness. To do this, we used RF-based models, which can better capture non-linear feature interactions (i.e., gene-gene interactions) (71,80) than linear models, to obtain predicted genetic interactions using SHAP (71). Models were trained with either individual or combinations of genetic variant types to identify gene-gene interactions. We focused on YPD Benomyl 500 µg/ml, because experimentally validated genetic interaction data are available for this environment. Models were trained using only benchmark benomyl genes from SGD (**S10 Table**). To assess the biological relevance of SHAP-based predictions, we compared them with experimentally validated genetic interaction networks from the BioGRID database and two additional publicly available networks: for benomyl 30 μg/ml (81) and a control condition (81) (**S11 Fig**). The SNP model performed similarly to the combined SNP + PAV + CNV, SNP + PAV, and SNP + CNV RF models (all models: testing R^2^ ≈ 0.55, **Fig 6A**). In contrast, combining PAVs and CNVs improved performance (testing R^2^ = 0.17) compared with PAVs alone (testing R^2^ = -0.09), but not over CNVs alone (testing R^2^ = 0.18). Model performances were not significantly associated with the number of features used to train the models (**S16 Table**). These results suggest that PAVs and CNVs of benomyl benchmark genes may not be as informative as SNPs in explaining fitness in the YPD Benomyl 500 µg/ml environment and that SNPs may be able to identify more relevant genetic interactions than other variant types in this environment.

**Fig 6.**
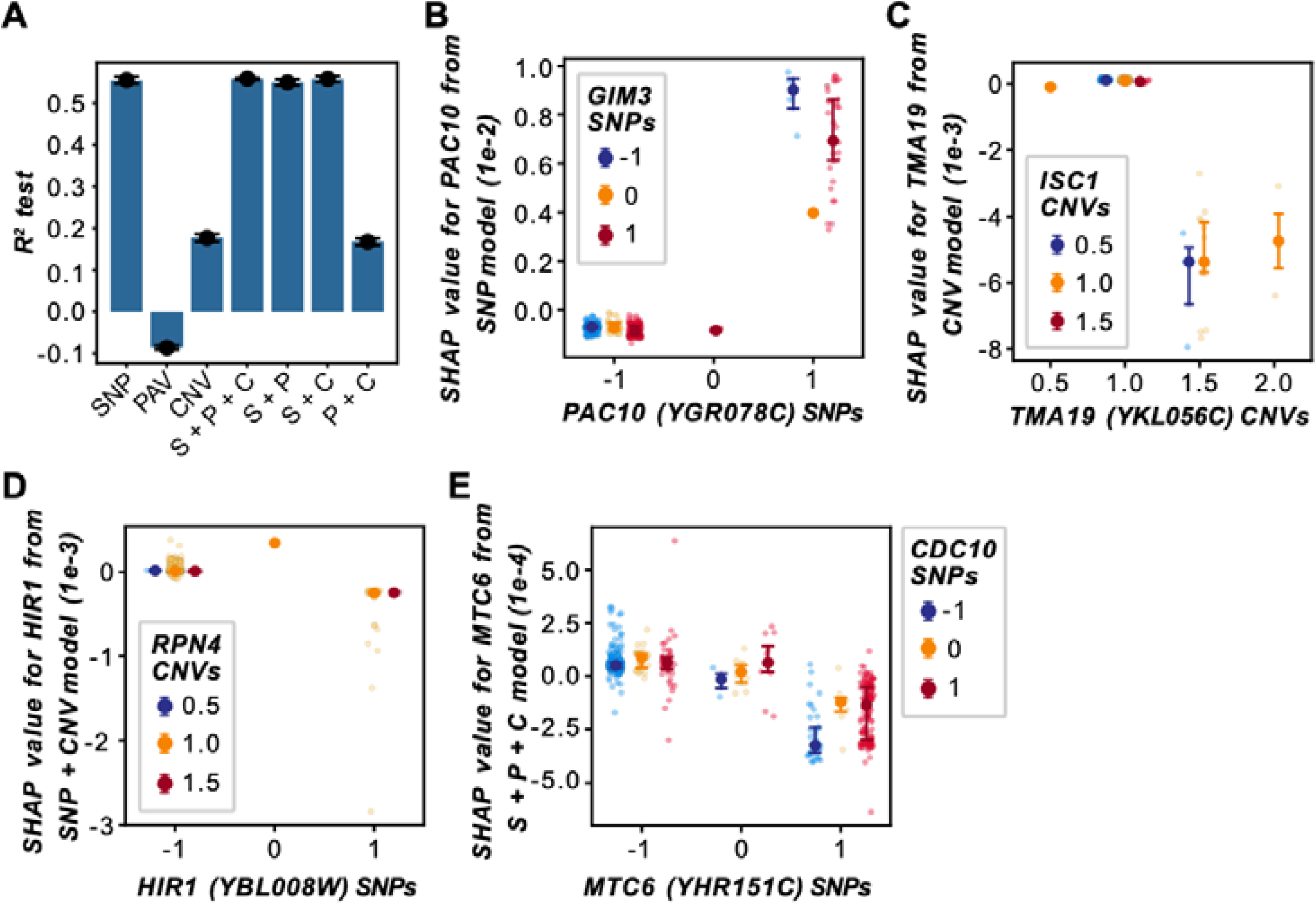
Contribution of genetic interactions to fitness predictions. **(A)** Test set performance R^2^ values for the prediction of fitness in YPD Benomyl 500 μg/ml by RF models trained on the benomyl benchmark genes. SNP, PAV, and CNV variants were mapped to genes (see **Materials and Methods**), and one variant with the highest feature importance was selected to represent the gene in the model. Models were trained on individual SNP, PAV, and CNV datasets or integrated datasets (S: SNP, P: PAV, C: CNV). **(B–E)** Feature interactions between **(B)** *PAC10* SNP and *GIM3* SNP, **(C)** *TMA19* CNV and *ISC1* CNV, **(D)** *HIR1* SNP and *RPN4* CNV, and **(E)** *MTC6* SNP and *CDC10* SNP. Axes: SNP or CNV genotypes of a gene (x-axis) and the gene’s corresponding SHAP values (y-axis). SNP genotypes are encoded as -1 (homozygous for the major allele), 0 (heterozygous), or 1 (homozygous for the minor allele). Points represent isolates and the color represents the SNP, PAV, or CNV genotypes of the second gene. Points with error bars represent the median SHAP value and the interquartile range at the 25th and 75th percentiles.

We identified a total of 210,758 feature interactions from the YPD Benomyl 500 µg/ml RF models in **Fig 6A**, excluding the PAV model because of its negative performance (**S17 Table**); of these 69,486 represented unique gene-gene interactions. We reasoned that if these gene-gene interactions are biologically meaningful, they may be identified by multiple genetic variant types. To examine this, we determined which types of variant-variant feature interactions mapped to the same gene pairs (**S18 Table**), and whether those gene pairs have been experimentally validated according to the literature. Consistent with our hypothesis, there was substantial overlap in the gene-gene interactions identified by different variant combinations (**S12 Fig**). In addition, although only one of the unique gene-gene interactions overlapped with the Benomyl 30 μg/ml network, 59 overlapped with the control condition network and 6,358 overlapped with BioGRID interactions. Of the 6,358 validated genetic interactions, 5,645 exhibited feature interactions for two or more variant-variant pair types (**S18 Table**), with SNP-SNP interactions identifying the most validated genetic interactions (6,095 out of 6,358 gene pairs). Similar to what we discovered for the benchmark genes, these experimentally validated genetic interactions were not significantly enriched in the SHAP-based feature interactions we identified (*q* > 0.38, **S19 Table**) regardless of the rank percentile examined (see **Materials and Methods**). However, the SHAP-based interactions may represent novel, biologically relevant genetic interactions that are good candidates for experimental validation. Furthermore, SHAP-based feature interactions provide insight into potential genetic mechanisms affecting the ability of interacting genes to contribute to fitness predictions. For example, the single Benomyl 30 μg/ml genetic interaction was between *PAC10* (Perish in the Absence of Cin8p, *YGR078C*) and *GIM3* (Gene Involved in Microtubule biogenesis, *YNL153C*, ranked 21, **S17 Table**). Both genes encode components of the prefoldin co-chaperone complex, which promotes α- and γ-tubulin formation (82). The minor allele of the *PAC10* SNP contributes to higher predicted fitness when it is homozygous and the *GIM3* SNP is heterozygous, and even higher predicted fitness when the *GIM3* SNP is homozygous for either the major or minor allele (**Fig 6B**).

We next examined the top five feature interactions with the highest SHAP interaction values from each model (excluding the PAV model). Of these 30 interactions, five were CNV-CNV interactions from the CNV model involving *TMA19* (Translation machinery associated, *YKL056C*). *TMA19* interacted with *ACL1* (Ankyrin repeat chaperone of Rpl1p, *YCR051W*, ranked 1st), *YCL001W-A (*an uncharacterized ORF, ranked 2nd), *IMP21* (Independent of mitochondrial particle, *YIL154C*, ranked 3rd), *ISC1* (Inositol phosphosphingolipid phospholipase C, *YER019W*, ranked 4th), and *YCR025C* (an uncharacterized ORF, ranked 5th, **S17 Table**). Although no genetic or physical interactions have been reported in the literature between these genes, the feature interaction between *TMA19* and *ISC1* makes biological sense given the functions of Tma19p and Isc1p. Tma19p translocates to the outer membrane of the mitochondria under stress conditions, including benomyl treatment, as an anti-apoptotic measure (83) and binds with microtubules to stabilize them (83). Isc1p also translocates to the mitochondria (84), modulates apoptosis by generating bioactive ceramide molecules (85), and is implicated in spindle elongation (86). The involvement of these genes in apoptosis and cell division may suggest genetic interactions or functional associations, and they are good targets for further experimental validation. Another interesting finding is the association between the copy numbers of *TMA19* and those of *ISC1*. For example, when specific isolates have one copy of *ISC1* and 1.5 (i.e., a full copy and an additional partial copy of the full length of the ORF) or 2 copies of *TMA19*, the *TMA19* CNV tends to contribute to lower predicted fitness (**Fig 6C**).

Among the five strongest feature interactions in the SNP + CNV model, two were SNP-CNV interactions of *HIR1* (Histone regulation, *YBL008W*). *HIR1* interacted with *SER1* (3-Phosphoserine aminotransferase, *YOR184W*, ranked 1st) and *RPN4* (Regulatory particle non-ATPase, *YDL020C*, ranked 3rd), the latter of which has been reported by multiple high-throughput studies to negatively interact with *HIR1* (87–89). Here we found that the contribution of the *HIR1* SNP to fitness predictions depends on the *RPN4* copy number in certain isolates. In isolates with a single copy or partial duplication of *RPN4* and a homozygous SNP genotype for the minor allele of *HIR1*, the *HIR1* SNP contributed to lower predicted fitness (**Fig 6D**).

Of the top 30 feature interactions, 14 were SNP-SNP interactions from the SNP, SNP + PAV, and SNP + PAV + CNV models. In the SNP + PAV + CNV model, SNP-SNP interactions often involved *CDC10* (Cell division cycle septin, *YCR002C*), which interacted with *JAC1* (J-type accessory chaperone, *YGL018C*, ranked 1st), *MTC6* (Maintenance of telomere capping, *YHR151C*, ranked 2nd), and *ISC1* (ranked 4th). Costanzo *et al.* (90) confirmed a negative genetic interaction between *MTC2*, another protein involved in maintenance of telomere capping, and *CDC10* under standard growth conditions; thus, *MTC6* may also genetically interact with *CDC10*. Examining the *MTC6* and *CDC10* feature interaction showed that regardless of the genotype of the *CDC10* SNP, the *MTC6* SNP contributes to reduced predicted fitness when isolates are homozygous for the minor allele and to higher predicted fitness when isolates are homozygous for the major allele (**Fig 6E**).

Overall, SHAP interaction values are useful for identifying gene pairs that are jointly important for predicting fitness. While enrichment of experimentally validated genetic interactions was not observed, several high-ranking feature interactions involved genes with reported genetic interactions, underscoring the value of utilizing SHAP interaction values for identifying candidate genes for functional validation. The lack of enrichment of experimentally validated genetic interactions may reflect the dependence of genetic interactions on the genetic background and the environment (81). For example, a subset of experimentally validated genetic interactions were obtained under a less severe benomyl stress (30 μg/ml) than the condition in which the isolates were grown (benomyl 500 μg/ml and YPD media). Furthermore, the experimentally validated genetic interactions also involved non-benomyl benchmark genes, which may also partially explain the lack of enrichment. In addition, several of the top 30 SHAP interactions were observed among genes with related functions, which may be indicative of genetic interactions (81,90,91), but further experimental validation is required for confirmation.

## Conclusion

Our study demonstrates that the type of genetic variant—SNP, PAV, or CNV—has a significant impact on both the accuracy of fitness predictions and the biological interpretability of predictive models. Furthermore, structural variants, particularly CNVs, contributed more strongly to predictive performance in most environments and were effective in identifying both known benchmark genes and novel candidate genes involved in environmental stress responses in the environments we assessed. One reason for this is the substantial effects that structural variants such as CNVs can have on gene expression and protein abundance, which in turn lead to fitness variation (62,72). For example, CNVs may decrease fitness when they are associated with increased abundance of protein, leading to elevated intracellular solute concentrations, which causes hypo-osmotic stress (62,92). On the other hand, CNVs can provide a temporary fitness advantage if they result in increased expression of a gene required for survival in an environment, but decrease fitness when the stress is removed (62). Furthermore, the number of CNVs may differ from that of SNPs or other variants depending on the genetic backgrounds or fungal species investigated (62,72). Thus, future studies of the genotype-phenotype relationship should strongly consider leveraging CNVs to better understand phenotypic variation and adaptation.

Importantly, our models were able to recover key benchmark genes in multiple environments—caffeine, benomyl, CuSO_4_, and sodium meta-arsenite—and candidate non-benchmark genes driving improvements in model performance, suggesting that non-benchmark genes explain a substantial portion of fitness variation. We also found that the genetic background of isolates influenced how individual genes contributed to fitness, with background-dependent effects being driven by many genes instead of a few high-impact genes for five out of 35 environments. Further studies on the effect of genetic background on fitness variation and how the environment alters the contributions of genotypes to fitness are needed to disentangle this complicated relationship.

Overall, our results highlight the complex interplay between genetic variation, environment, and genetic background in shaping fitness. By identifying both shared and environment-specific candidate genes, this study provides insights into the genetic basis of fitness variation in different environments and a foundation for future functional validation experiments. The comparative analysis of benchmark and candidate genes across environments using SHAP values reveals common and distinct mechanisms of fitness, while clustering of isolate-level gene contributions uncovers patterns of coordinated or opposing effects across genetic backgrounds. Together, these findings offer a framework for guiding gene target selection in engineering yeast strains with improved stress resilience.

## Materials and Methods

### Data pre-processing, kinship calculation, and estimation of population structure

Three types of genetic variant data—single nucleotide polymorphisms (SNPs), open reading frames (ORFs) of presence/absence variants (PAVs), and ORF copy number variants (CNVs)—and fitness measurements (**S1 File**) for 750 diploid *S. cerevisiae* isolates grown in 35 different environments were obtained from (51). Here, fitness is the ratio between the size of colonies grown in an environment and that in the reference environment (YPD medium at 30°C). The SNP data were filtered using VCFtools v0.1.16 (93) to retain only biallelic SNPs with a minor allele frequency (MAF) > 5% and missing data < 20%, using the parameters “-- maf 0.05”, “--max-alleles 2”, “--min-alleles 2”, “--max-missing 0.2”, “--recode”, and “-- remove-indels”. The final genotype dataset used in this study included 118,382 SNPs (**S2 File**). Genotypes were re-coded into fastPHASE format with PLINK v1.9 (94,95) using the parameter “--recode12 fastphase”. Missing genotypes were imputed using fastPHASE v1.4.8 with the parameter “-T10” (96). Genotypes were encoded as {-1, 0, 1} corresponding to {AA, Aa, aa}, where A is the major allele and a is the minor allele. Kinship between isolates was calculated using the centered identity-by-state method (97) implemented in TASSEL v5 (98) with the parameters “-KinshipPlugin” and “-method Centered_IBS” (**S3 File**). Population structure was modeled as the first five principal components (PCs) of the SNP genotypes estimated with the Scikit-learn v1.2.2 (99) Principal Component Analysis function (**S4 File**). Eighty-eight ORFs with missing PAV and CNV values in all isolates were excluded, resulting in PAV and CNV datasets with 7,708 ORFs out of the original 7,796 reported by (51) (**Files S5** and **S6,** respectively).

### Predictive modeling of fitness in each environment using genomic prediction

For each of the 35 environments, a “single-environment” model was built where the complete set of SNP, PAV, CNV, or PC values (referred to as features) was used to predict fitness with linear and non-linear methods. Three linear models were implemented in R v4.3.2: Ridge Regression Best Linear Unbiased Predictor (rrBLUP) using rrBLUP v4.6.3 (100), Bayesian-Least Absolute Shrinkage and Selection Operator (Bayesian LASSO) (101), and BayesC (102). Bayesian LASSO and BayesC were implemented using BGLR v1.1.4 (103) with 32,000 iterations, where the first 3,200 iterations were discarded as burn-in. Non-linear, machine learning regression models were implemented in Python 3.11.5 using Scikit-learn v1.2.2 for Random Forest (RF) (104) and XGBoost v2.0.3 (105) for eXtreme Gradient Boosting (XGBoost) (105).

Before building any model, one-sixth of the yeast isolates were randomly held out as the test set, which was used exclusively to evaluate model performance. The remaining five-sixths of the isolates, referred to as the training set, were used to train the models. For rrBLUP models, the rrBLUP R package automatically estimates the regularization and kernel parameters from the data, so no hyperparameter tuning was conducted with cross-validation. No hyperparameter tuning was conducted for BayesC or BayesianLASSO either. rrBLUP, BayesC, and Bayesian LASSO training were conducted within a five-fold cross-validation scheme and repeated 20 times. For machine learning algorithms, hyperparameter tuning was conducted within a five-fold cross-validation scheme, which was repeated 10 times for RF and 100 times for XGB, to account for the larger hyperparameter space in XGB. The hyperparameters for RF, which were tuned using Scikit-Learn’s GridSearchCV function, were “max_depth” [3, 5, 10], “max_features” [0.1, 0.5, “sqrt”, “log2”, “None”], and “n_estimators” [100, 500, 1000]. The hyperparameters for XGBoost, which were tuned using HyperOpt v0.2.7 (106), were "learning_rate" [0.01 to 0.4], "max_depth" [2 to 10] with a step size of 1, "subsample" [0.5 to 1.0], "colsample_bytree" [0.7 to 1], and "n_estimators" [5 to 500] with a step size of 5. To choose the best parameter combinations, we used the negative mean squared error for RF in GridSearchCV and the negative mean R^2^ for XGBoost in HyperOpt. The best combination of parameters was chosen based on the validation set performance and used to train a new model within a five-fold cross-validation scheme. The coefficient of determination (R^2^) was calculated between the observed and predicted relative fitness values, and the average of the 20 training repetitions was used as the validation set performance of RF and XGB.

### Feature importance and feature selection

The impact of features on the performance of the RF models was assessed using two methods. The first was Gini importance (70). For SNPs, the top {2*^n^* | *n* ∈ L, 1 ≤ *n* ≤ 10} □ {1000 × *n* | *n* ∈ □, 1 ≤ *n* ≤ 30} features, where *n* is the number of features, were selected based on average Gini importance values, which we refer to as “Gini importance”. The Gini importance was calculated by taking the average feature importance across the 20 training repetitions of the RF models built using the complete feature sets. These feature subsets were used to build new RF models to find the minimum feature set size required to reach peak training performance (referred to as an optimized RF model). For PAV and CNV features, the top {2*^n^* | *n* ∈ □, 1 ≤ *n* ≤ 10} □ {250 × *n* | *n* ∈ □, 1 ≤ *n* ≤ 30} features were selected. The optimized single-environment RF models were selected based on the inflection point of a feature selection curve (x-axis: number of features, y-axis: average performance R^2^ across 10 training iterations on the validation set). These feature subsets were then used to build models for all other algorithms. Gini importance values for SNP, PAV, and CNV features for RF models using the complete feature sets can be found in **S7 File**.

SHAP v.0.42.1 (39,71) was also used to assess feature importance. SHAP values for each feature were estimated for each isolate used for training RF models on the complete or optimized feature sets. Because training was repeated 20 times for RF models built using complete feature sets and 10 times for optimized RF models, the model with the highest validation performance, which was assumed to represent the data the best, was used to estimate SHAP values. The SHAP values for SNPs, PAVs, and CNVs from the RF models trained on the complete feature sets are provided in **S7 File**.

### Predicting model performances using technical and fitness-related features

The effects of the features narrow-sense heritability (*h^2^*) of fitness, median fitness (med), and fitness variance (var) on performance (y) of the 35 PC, SNP, PAV, or CNV optimized single-environment RF models were estimated by ordinary least squares (OLS) regression using the statsmodels v0.14.2 Python package. Feature effects were estimated using the following equation:

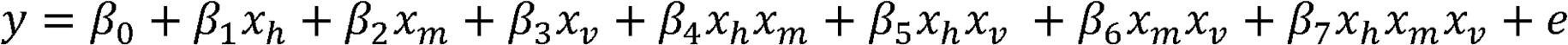

where *β* is a vector of coefficients and *β*_0_ is the intercept; *x* is a vector of values of the features *h^2^* (*h*), med (*m*), and var (*v*); and e is a vector of residuals. An additional OLS regression model was built using the number of features used to train the optimized RF models. Regression models were built to assess the performance of the optimized RF models built with one genetic variant type (i.e., PCs, SNPs, PAVs, or CNVs). Linear model goodness of fit was evaluated using both the R^2^ and adjusted R^2^, where the latter corrects the R^2^ for the number of predictor variables relative to the sample size to prevent overestimating model fit (107). SHAP values were estimated for each linear model using the LinearExplainer function to further assess feature effects on model performances.

The *h^2^* of fitness in all 35 environments was estimated using a mixed model equation implemented in the Sommer v4.3.0 R package (108) using the mmer function. The formula for the mixed model was as follows:

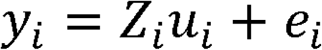

where *y_i_* is a vector of fitness values, *u_i_* is a vector of random effects, *e*_i_ are residuals for environment *i* (*i* = 1, …, 35), and *Z_i_* is an incidence matrix of random effects. A 750 × 1 vector of trait values for environment *i* was used to represent *y_i_*. A 750 × 750 additive relationship matrix and a 750 × 750 dominance relationship matrix were estimated using the SNP genotypes and used as random effects.

### Mapping of SNPs and ORFs to genes

The *S. cerevisiae* reference genome (S288C; version R64-3-1) and gene annotation files (gff3) were obtained from the *Saccharomyces* Genome Database (SGD; http://sgd-archive.yeastgenome.org/sequence/S288C_reference/) and used to obtain the list of S288C genes and their translational start and stop positions. SNP variants were assigned to genes if they were found in genic regions (**S8 File)**. ORFs comprising PAV and CNV features were assigned to S288C genes based on reciprocal best match in two steps. In the first step, sequence similarity between query ORF nucleotide sequences and the protein sequences of S288C was determined using BLASTx (109,110) with parameters “-max_target_seqs 2”, “-max_hsps 1”, and “--evalue 1e-06”. The top matching S288C gene, G, was considered a candidate gene to a query ORF, O, if the percent identity was ≥ 95% and the E-value was < 1e-6.

In the second step, ORF-to-gene mapping was finalized by using the nucleotide sequence of the candidate S288C gene, G, identified in the first step as the query to search against non-S288C ORF nucleotide sequences using tBLASTx with the same parameters as the first step. If the top match of an S288C gene G remained the ORF O, based on the same filter criteria in the first step, O was mapped to G. Out of 7,708 ORFs, 5,902 mapped to 5,873 unique S288C genes, and 16 of these ORFs mapping to > 1 gene were excluded. To obtain systematic gene names for the mapped genes, identifiers of genes from the non-redundant database were mapped to the S288C gene identifiers using the NCBI Datasets Gene tool (https://www.ncbi.nlm.nih.gov/datasets/gene/) and the SGD YeastMine Gene List tool (https://yeastmine.yeastgenome.org/yeastmine/bag.do). The ORF-to-gene mapping data can be found in **S9 File**.

### Feature rank percentile correlations

Spearman’s rank correlation between the Gini importance and the average absolute SHAP values of SNP, PAV, or CNV features from the optimized RF models was determined after dropping features with zero importance values. The remaining features were ranked using the Pandas “DataFrame.rank” function and the “average” method, and the correlation was calculated based on the features common to both importance measures. Similarly, for correlations based on the RF models trained on the complete feature sets, features with zero importance values were excluded, and the remaining features were ranked.

To assess relationships between SNP and PAV or CNV features from the optimized RF models, these features were mapped to genes and were ranked according to either the maximum Gini importance or the maximum average absolute SHAP value (calculated across the isolates) of all the features that mapped to the same genic region (see **Mapping of SNPs and ORFs to genes**). SNP, PAV, and CNV features that did not map to any gene, including intergenic SNPs and/or had a zero Gini importance or average absolute SHAP value, were excluded from the ranking. The Spearman’s rank correlation was calculated between SNPs vs PAVs or CNVs, and PAVs vs CNVs.

To assess relationships between environments from the optimized RF models, we determined the number of overlapping genes across the 35 environments. Features from RF models trained on the optimized SNP, PAV, or CNV feature sets were mapped to genes. To determine the number of environments in which a gene appeared within the optimized feature set, genes with non-zero feature importance—the highest average absolute SHAP value or the highest Gini importance of the features mapped to that gene—were assigned a value of 1, creating a gene presence matrix. These presence values were summed across all environments, and the resulting counts were compared to a null distribution of median counts generated from 10,000 permutations of the gene presence matrix in which each column (environment) in the gene presence matrix was permuted individually. The actual distribution of environment counts was compared to the randomized null distribution using the Kolmogorov-Smirnov test (alternative = “greater”).

Spearman’s rank correlation of shared features between two environments was estimated after dropping features with non-zero importance in at least one environment and ranking the remaining features for each environment. Correlations were determined for ranks based on Gini importance or average absolute SHAP values of shared features.

### Benchmark fitness genes and known genetic interactions

Candidate known fitness genes were obtained from SGD by searching for “benomyl”, “caffeine”, “copper(II) sulfate” (CuSO_4_), and “sodium arsenite” (also known as sodium meta-arsenite). Phenotype annotations of candidate genes found in the “Chemicals” category page for these compounds were filtered by the “Phenotype” and “Mutant Information” values based on the following criteria. A candidate gene was considered as a benchmark fitness gene for the target condition if it had mutant information annotations matching "null Allele", "reduction of function", or "reduction of function Allele" and phenotype annotations matching "resistance to chemicals: decreased", "viability: decreased", "metal resistance: decreased", "oxidative stress resistance: decreased", "respiratory growth: decreased", or "stress resistance: decreased". After the filtering step, 386 genes with experimental evidence of an effect on fitness compared to a reference condition were identified for benomyl, 752 for caffeine, 162 for CuSO_4_, and 280 for sodium meta-arsenite. S288C genes with mapped SNP features included 370/386 benomyl fitness genes, 725/752 caffeine fitness genes, 156/162 CuSO_4_ fitness genes, and 278/280 sodium meta-arsenite fitness genes. The ORF features mapped to 350/386 benomyl fitness genes, 688/752 caffeine fitness genes, 145/162 CuSO_4_ fitness genes, and 270/280 sodium meta-arsenite fitness genes. In addition, a list of manually curated genes for benomyl (5 genes), caffeine (10 genes), CuSO_4_ (15 genes), and sodium meta-arsenite (15 genes) were obtained from the literature.

Experimentally validated genetic interaction information for S288C was collected from the BioGRID database (https://thebiogrid.org/). Gene pairs with the evidence annotations "Synthetic Growth Defect", "Synthetic Lethality", "Synthetic Rescue", "Negative Genetic", and "Positive Genetic" were selected, totaling 438,546 genetic interactions. Additional genetic interactions that were experimentally verified using single and double mutant yeast strains grown under a control condition (glucose) or in 30 µg/ml benomyl were obtained from data published by Costanzo et al. (81). These genetic interactions observed under control conditions or in the presence of benomyl were filtered according to a stringent confidence threshold (*p* < 0.05 and |ε| > 0.12, where ε is the genetic interaction score), yielding 3,417 genetic interactions in the control condition and 3,472 genetic interactions in the presence of benomyl. The combined set of unique genetic interactions (including BioGRID and (81) data) resulted in 441,520 gene pairs (**S10 File**).

### Gene ontology term, metabolic pathway, benchmark gene, and experimentally validated genetic interaction enrichment analyses

Gene ontology (GO) term annotations v.20220912 were retrieved from the GO GAF v2.2 format (sgd.gaf; http://current.geneontology.org/annotations/index.html). GO terms with experimental evidence codes (IDA, IPI, IMP, IGI, IEP, HDA, HMP, HGI, HEP) were kept and mapped to the features. Pathway annotations (downloaded October 10, 2022) were retrieved from the MetaCyc database (https://metacyc.org/group?id=biocyc14-55140-3843260367). Before conducting enrichment analyses for GO terms and pathways, SNP, PAV, and CNV features were mapped to the S288C genes as detailed in **Mapping of SNPs and ORFs to genes**. Gene features that met the feature selection cut-off criteria were referred to as “important genes” for predicting fitness in an environment, i.e., these features were used to train the optimized RF models. For features that did not meet the feature selection cut-off, the genes they mapped to were considered as the background (unimportant) gene set. Features that did not map to any genes or mapped to multiple genes were excluded from the analysis.

Enrichment of important genes for a GO term or a pathway annotation, *A*, was determined by calculating four values in a 2 × 2 contingency table: *a —* the number of important genes with *A*, *b —* the number of background genes with *A*, *c —* the number of important genes without *A*, and *d —* the number of background genes without *A*. For each annotation, these four values were used to determine the enrichment *p* with two-sided Fisher’s exact tests. To correct for multiple testing, hypothesis tests with *P =* 1 were removed, and the remaining *p-*values were corrected using the Benjamini and Hochberg method (111).

For analysis of benchmark fitness gene enrichment in each environment, we tested whether benchmark genes related to that environment (i.e., benomyl benchmark genes were identified within the YPD Benomyl 500 μg/ml model) or to a different environment (e.g., benomyl benchmark genes were identified within the YPD Caffeine 40 mM model) were enriched at different rank percentile thresholds (within the top 1%, 5%, 10%, 15%, 20%, or 25% of ranked genes) of Gini importance or average absolute SHAP values for each optimized model. Benchmark genes associated with benomyl, caffeine, copper(II) sulfate, and sodium meta-arsenite stress were sourced from both SGD and the literature (see **Benchmark fitness genes and known genetic interactions**). Enrichment was assessed among genes within the top 1%, 5%, 10%, 15%, 20%, or 25% rank percentile based on SNP, PAV, or CNV features. Features were mapped to genes, and rank percentiles were calculated using either the highest Gini importance or the highest average absolute SHAP value of the features mapped to that gene. Features that had non-zero importance, did not map to a gene, or mapped to multiple genes were excluded from the rankings. To ensure that all genes in the yeast genome represented by the SNP, PAV, or CNV features were included in the analysis, gene rankings from the optimized single-environment RF models were combined with rankings of non-overlapping genes from models trained on the complete feature sets. Enrichment analysis was conducted for five environments (YPD Caffeine 40 mM, YPD Caffeine 50 mM, YPD Benomyl 500 μg/ml, YPD CuSO_ 10 mM, and YPD Sodium meta-arsenite 2.5 mM) in a similar manner to the GO term and pathway enrichment analyses. However, the annotation *A* represents whether the gene is a benchmark fitness gene in either the benomyl, caffeine, copper(II) sulfate, sodium meta-arsenite, or literature-based gene lists.

### Assessing the contribution of benchmark genes to fitness predictions

To assess the importance of benchmark genes for fitness predictions, we trained 45 new RF models for predicting fitness using reduced SNP, PAV, or CNV feature sets for five environments: YPD Caffeine 40 mM, YPD Caffeine 50 mM, YPD Benomyl 500 μg/ml, YPD CuSO_4_ 10 mM, and YPD Sodium meta-arsenite 2.5 mM. Three kinds of reduced feature sets were generated: (1) a feature set containing important non-benchmark genes (identified by the optimized single-environment RF models) plus benchmark genes (from the RF models trained on the original complete feature sets), (2) only important non-benchmark genes, and (3) only benchmark genes. SNP, PAV, or CNV features from the RF models built using complete feature sets and the optimized RF models were mapped to genes (see **Mapping of SNPs and ORFs to genes**). Features were obtained by representing each gene by the SNP within the genic region, PAV, or CNV feature with the highest average absolute SHAP importance. Then, benchmark and non-benchmark genes were determined based on their presence in the SGD benchmark gene lists. Intergenic SNPs were excluded. RF models were trained using Scikit-learn v1.2.2 and evaluated as described under **Predictive modeling of fitness in each environment using genomic prediction**. The number of features and validation and testing performances for these models are provided in **S11 File**.

### Analyzing genetic distance to S288C and the effect on benchmark gene importance

Genetic distances (Euclidean distances) between the 625 diploid isolates from the training set and S288C were estimated from SNP and PAV values. SNP genotypes were recoded to {0, 1, 2} genotype encodings, where 0 refers to a genotype homozygous for the reference allele, 1 for heterozygous, and 2 for homozygous for the alternative allele (**S12 File**). S288C SNP genotypes were denoted as a vector of 0s, since S288C is the reference genome and was not one of the training isolates. S288C PAV values were determined from the BLASTx and reciprocal tBLASTx results outlined in **Mapping of SNPs and ORFs to genes**, where an ORF was given a 0 value if it did not map to the S288C genome, or a 1 if it did. Euclidean distance between pairs of isolates was estimated using the scipy.spatial.distance.pdist function. No scaling was applied to the SNP or PAV matrices prior to calculating Euclidean distances. The distance matrices are provided in **Files S13** and **S14** for SNPs and PAVs, respectively.

To compare the maximum absolute SHAP values of benchmark genes between two clusters of isolates, K-means clustering was performed on the SNP and PAV genetic distance matrices using the KMeans function from Scikit-learn v1.2.2. For each distance matrix, the number of clusters (k) was varied from 2 to 10, and elbow plots were used to select the optimal number of clusters (k = 6 for SNPs, k = 4 for PAVs) based on inertia. No scaling was applied before clustering. The cluster containing S288C was identified, and the remaining clusters represent isolates that are the least genetically related to S288C. A two-sided Mann-Whitney U test (alternative = "greater") was used to compare the distributions of median absolute SHAP values of benchmark genes between these two clusters, using the Mann-Whitney U test as implemented in SciPy v1.11.4. To construct these distributions, SHAP values were obtained from either the optimized RF model or the complete RF model trained on all SNP or PAV features for a given environment. Only SHAP values corresponding to features that mapped to benchmark genes were retained. For each isolate in a cluster, the median absolute SHAP value was computed per gene.

### Clustering of SHAP values

Based on the median absolute SHAP values of the top 20 SNP, PAV, or CNV features from the optimized RF models, budding yeast isolates were clustered for five environments: YPD Caffeine 40 mM, YPD Caffeine 50 mM, YPD Benomyl 500 μg/ml, YPD CuSO_4_ 10 mM, and YPD Sodium meta-arsenite 2.5 mM. Hierarchical clustering was performed using the scipy.cluster.hierarchy.linkage function with the “ward” method and “euclidean” distance metric. To obtain clusters with different granularities, clusters were split by distance thresholds tailored to SNPs, PAVs, and CNVs for the YPD Caffeine 40 mM, YPD Caffeine 50 mM, YPD Benomyl 500 μg/ml, YPD CuSO_4_ 10 mM, and YPD Sodium meta-arsenite 2.5 mM environments (see **S15 File**). Features were also clustered for visualization purposes. SHAP values of the top 20 SNP, PAV, and CNV features for each model are provided in **S16 File**.

To assess the relationship between the SHAP values of isolates and fitness across clusters, the median absolute SHAP value across the top 20 features was calculated for each isolate in a cluster (*x* variable). Similarly, the median fitness of isolates was calculated for each cluster (*y* variable). A linear regression line was fitted between *x* and *y* using the scipy.stats.linregress function with default arguments. The function provided estimates of the slope, intercept, standard error of the intercept, Pearson correlation, and the *p*-value for a hypothesis test of whether the slope is zero (based on the Wald test using the *t*-distribution).

### Estimating SHAP interaction scores

To estimate SHAP interaction scores between feature pairs, three new RF models and three new rrBLUP models were built to predict fitness in YPD Benomyl 500 μg/ml using reduced SNP, PAV, or CNV feature sets. An additional four RF models were trained using integrated feature sets (SNP + PAV, SNP + CNV, PAV + CNV, or SNP + PAV + CNV). The reduced feature sets consisted of features that mapped to benomyl benchmark genes. The feature with the highest average absolute SHAP value was selected to represent each gene. Intergenic SNPs were excluded. These reduced SNP, PAV, and CNV feature sets were then concatenated column-wise to create the integrated feature sets. Of the 118,382 available SNP features, 370 were selected. Of the 7,708 PAV and CNV features, 350 were selected for each. RF models (implemented in Scikit-learn v1.5.2) and rrBLUP models were trained as described in **Predictive modeling of fitness in each environment using genomic prediction**. SHAP interaction scores were estimated from the RF model with the highest validation R^2^ among training repetitions for each reduced or integrated feature set. SHAP interaction scores between gene features were calculated using the shap.TreeExplainer.shap_interaction_values function (implemented in SHAP v.0.42.1).

## Acknowledgments

This work was supported by the National Science Foundation (DGE-1828149 and MCB-2218206 to S.-H.S. and K.S.A.; IOS-2107215 and MCB-2210431 to M.D.L. and S.-H.S.), the National Institute of General Medical Sciences of the National Institutes of Health under the Award Number T32GM110523 to K.S.A. This material is based upon work at the Great Lakes Bioenergy Research Center supported by the U.S. Department of Energy, Office of Science, Biological and Environmental Research under Award Number DE-SC0018409.

## Author contributions

K.S.A. and S.-H.S. conceptualized and designed the study; K.S.A. conducted the data processing and formal analysis; P.I. and G.d.l.C. assisted with the computational modeling. K.S.A. made the figures with help from P.I., G.d.l.C., M.L. and S.-H.S. K.S.A., P.I., G.d.l.C., M.L. and S.-H.S. wrote, reviewed, and edited the manuscript. All authors read and approved the final manuscript.

## Data availability

All data and code needed to reproduce the results from this study are available on Zenodo (https://doi.org/10.5281/zenodo.17245961) and GitHub (https://github.com/ShiuLab/Manuscript_Code/tree/master/2026_yeast_fitness_gxe).

## Supplemental figure legends

**S1 Fig.**
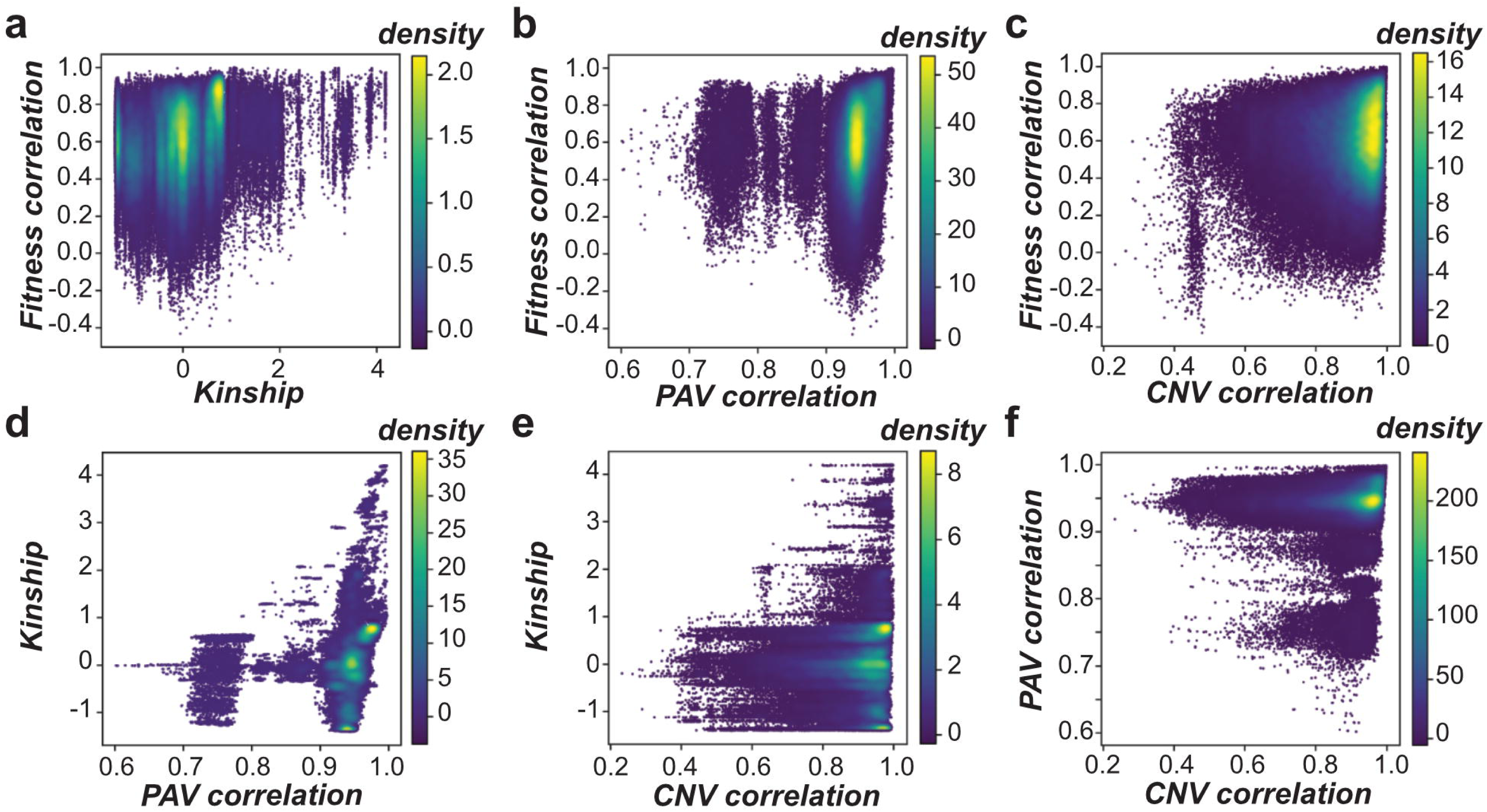
Correlations between fitness and genetic variant types and relationships between pairs of genetic variant types. Spearman’s rank coefficient (ρ) was calculated to compare the relationships between data type correlation matrices, which were calculated between pairs of isolates. **(A-F)** Density scatter plots showing the relationship between fitness correlation and kinship (ρ = 0.27, *p* < 2.2×10^-16^) **(A)**, fitness correlation and PAV correlation (ρ = 0.30, *p* < 2.2×10^-16^) **(B)**, fitness correlation and CNV correlation (ρ = 0.14, *p* < 2.2×10^-16^) **(C)**, kinship and PAV correlation (ρ = 0.48, *p* < 2.2×10^-16^) **(D)**, kinship and CNV correlation (ρ = 0.10, *p* < 2.2×10^-16^) **(E)**, and PAV correlation and CNV correlation (ρ = 0.17, *p* < 2.2×10^-16^) **(F)**.

**S2 Fig.**
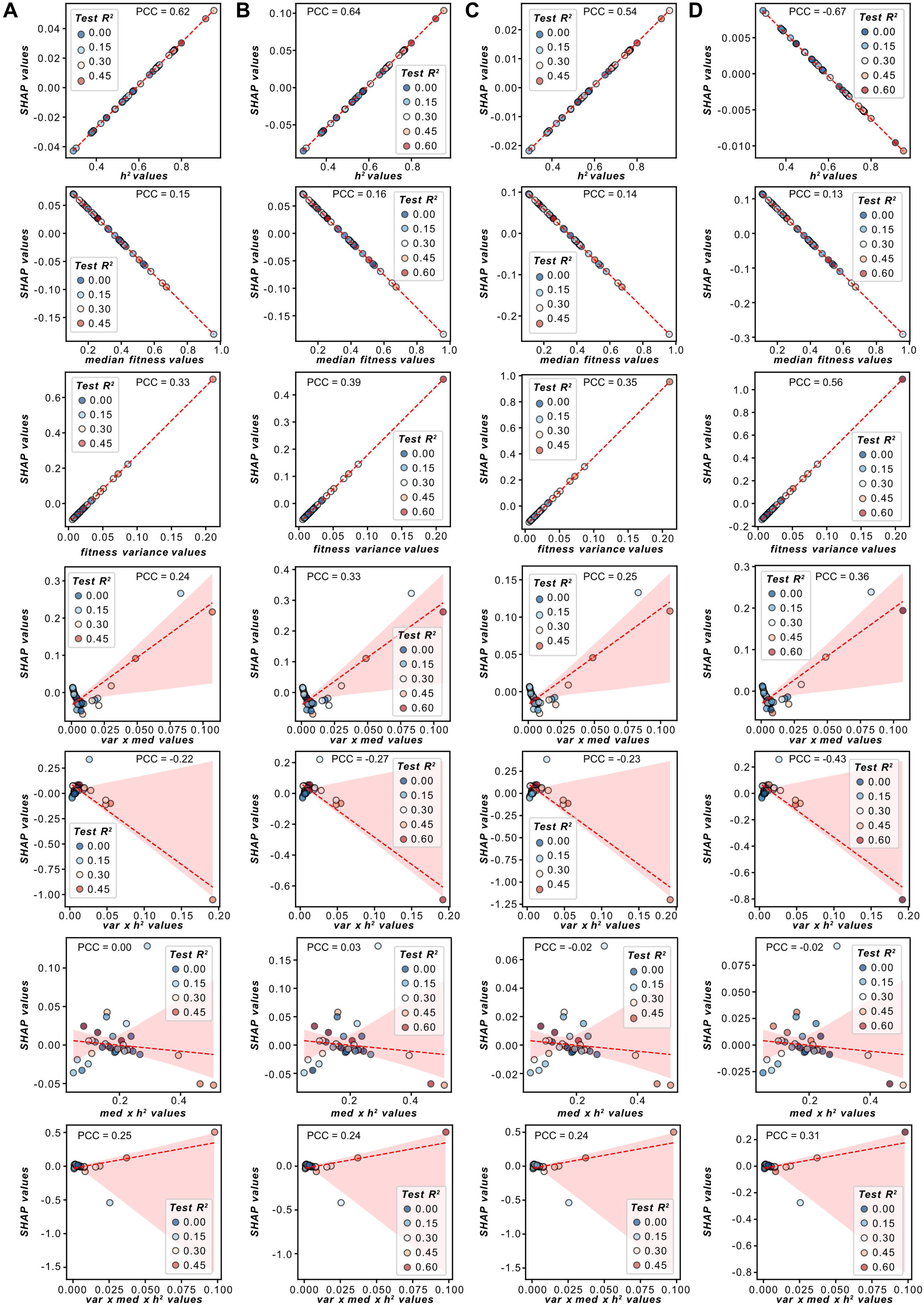
SHAP values of fitness-related features. **(A-D)** Scatterplots showing the relationship between the values of a fitness-related feature (x-axis, i.e., *h^2^*, median fitness, fitness variance, and their pairwise and three-way interactions) and the SHAP values of that feature (y-axis) from each **(A)** PC-, **(B)** SNP-, **(C)** PAV-, and **(D)** CNV-based linear model that was built to predict single-environment optimized RF model performances. Points are colored by the testing set R^2^ performance. The Pearson correlation values between the SHAP values of a feature versus the testing set R^2^ are shown.

**S3 Fig.**
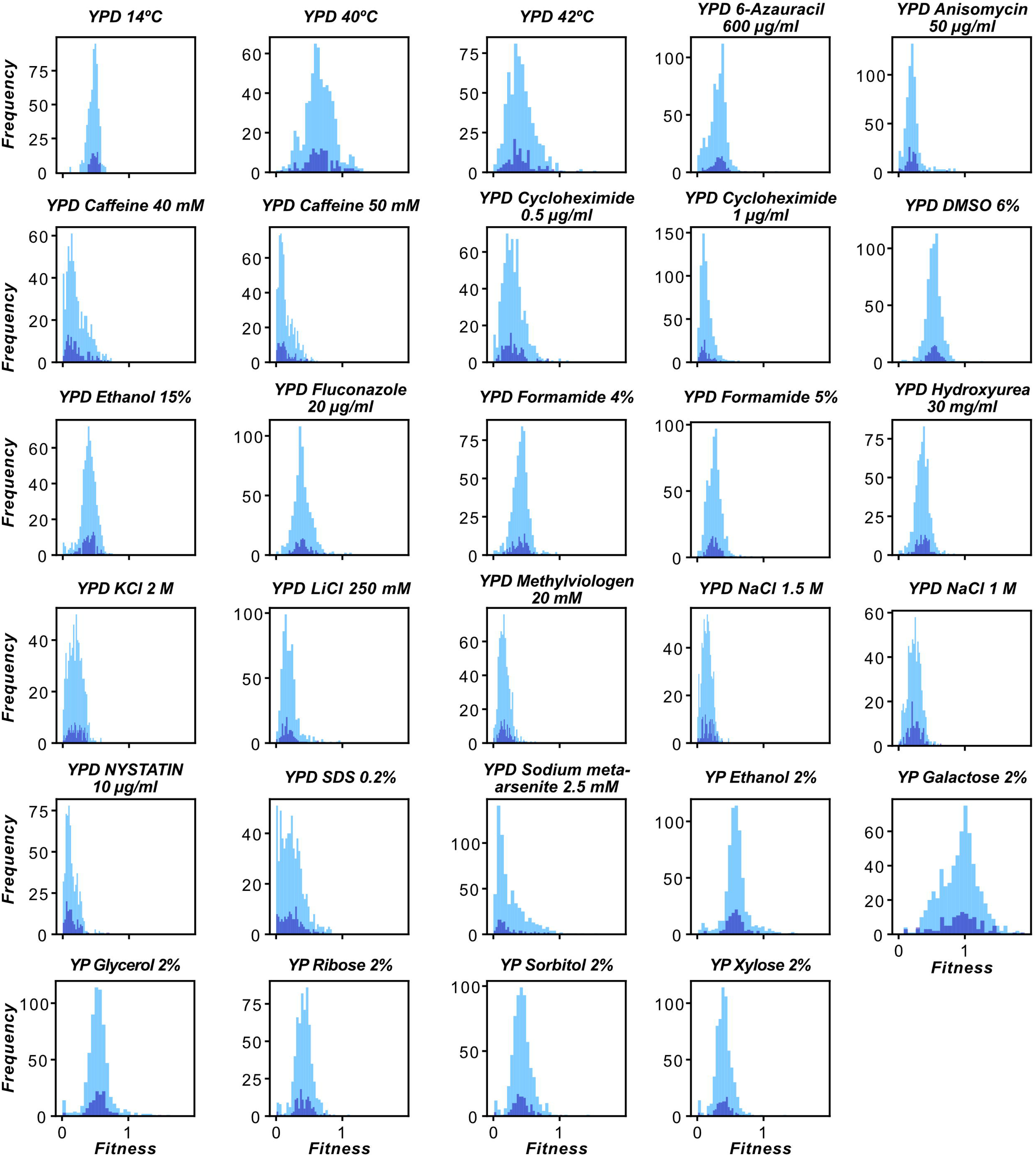
Fitness distributions of isolates in each environment. Fitness distributions of the 625 diploid isolates used for training models (blue) and the 125 isolates (purple) used for testing are shown for 29 environments.

**S4 Fig.**
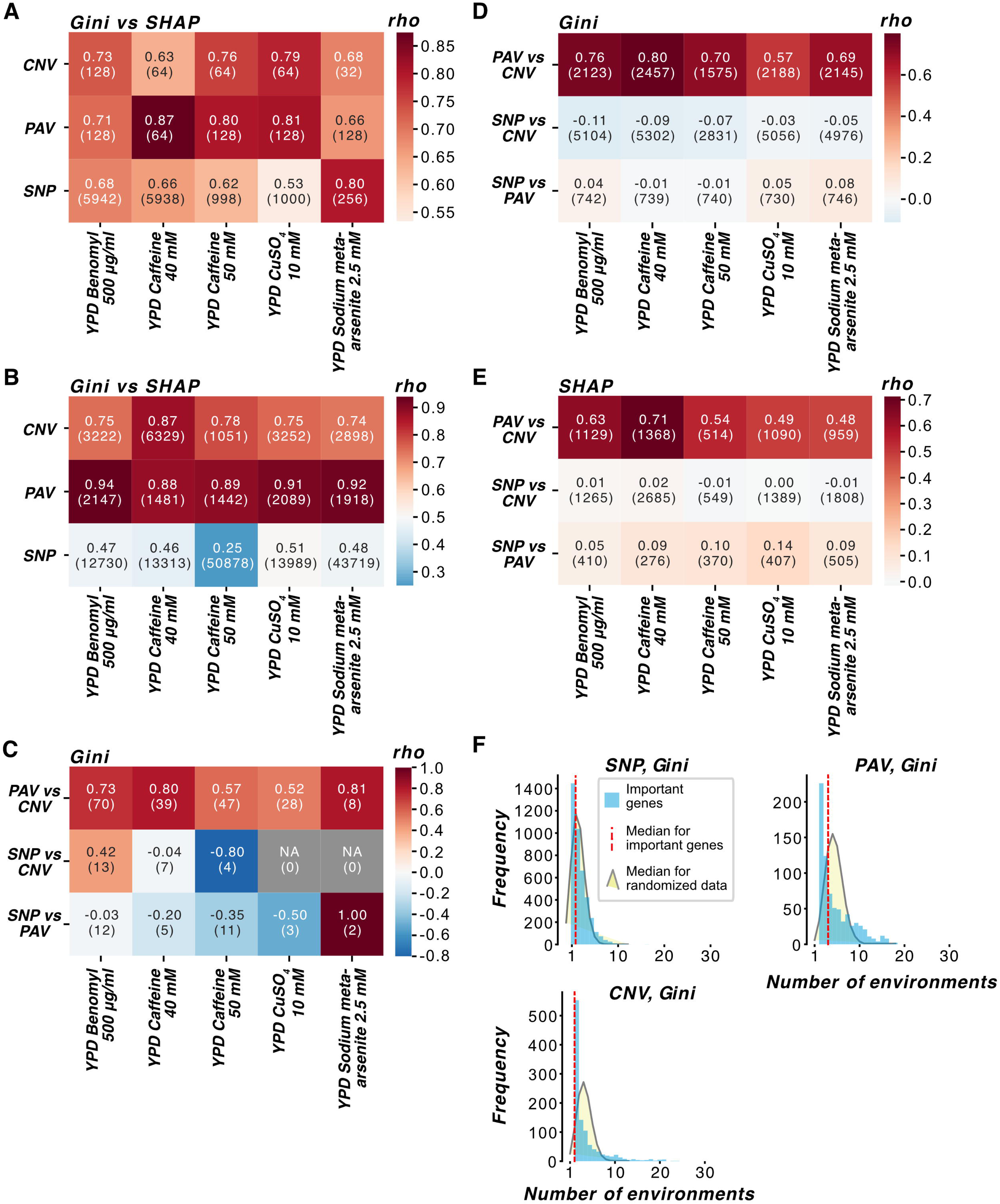
Relationships between Gini and SHAP feature importance in different environments. **(A-B)** Spearman’s rank correlation between feature rankings based on Gini importance and average SHAP values for **(A)** the optimized single-environment RF models and **(B)** single-environment RF models trained on complete SNP, PAV, or CNV feature sets. The number of overlapping features are shown in the parenthesis. **(C-E)** Spearman’s rank correlations between genetic variant types using Gini feature importance from the **(C)** optimized RF models and **(D)** the RF models built using complete feature sets and **(E)** using average SHAP values for the RF models trained on complete feature sets. The number of overlapping genes are shown in the parenthesis. See **Materials and Methods** for details about the gene to feature mappings. **(F)** Gini importance values from the optimized RF models were used to determine the distribution of unique or shared genes with non-zero importance across 1 to 35 environments (blue bars; red line: median). This distribution was compared to a null distribution of median randomized counts (see **Materials and Methods**) using the Kolmogorov-Smirnov test (alternative = “greater”); for SNPs: median *P* = 5.9×10^-28^; for PAVs: *P* = 1.5×10^-31^; for CNVs: *P* = 4.7×10^-71^.

**S5 Fig.**
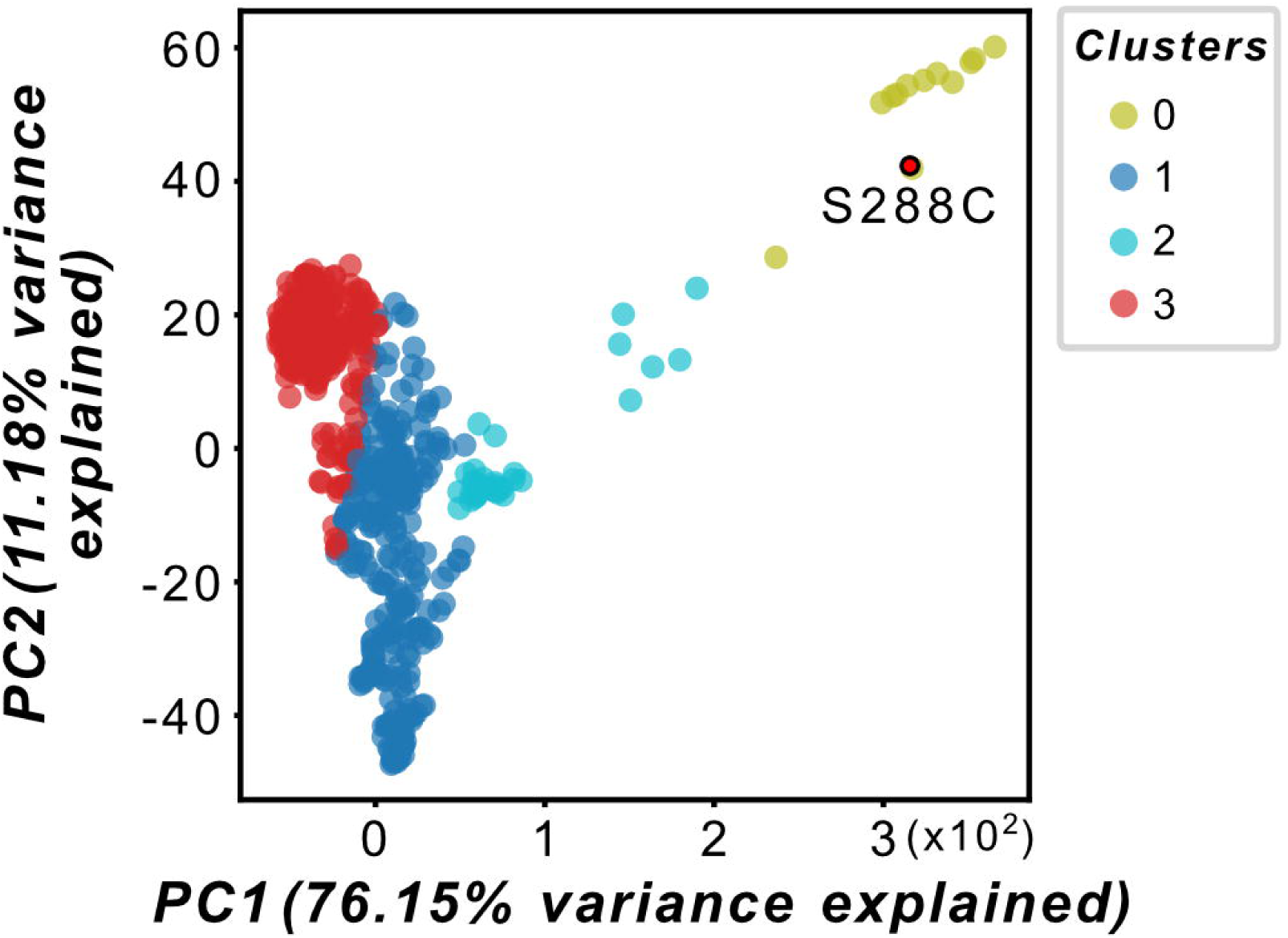
Genetic relatedness of isolates to laboratory strains based on PAVs. Principal component analysis of the Euclidean distance matrix of the PAV genotypes was performed to assess genetic relatedness among isolates. Clusters of genetically similar isolates were identified using K-means clustering applied to the distance matrix. Four clusters were identified from the PAV matrix. S288C is labeled in the plot.

**S6 Fig.**
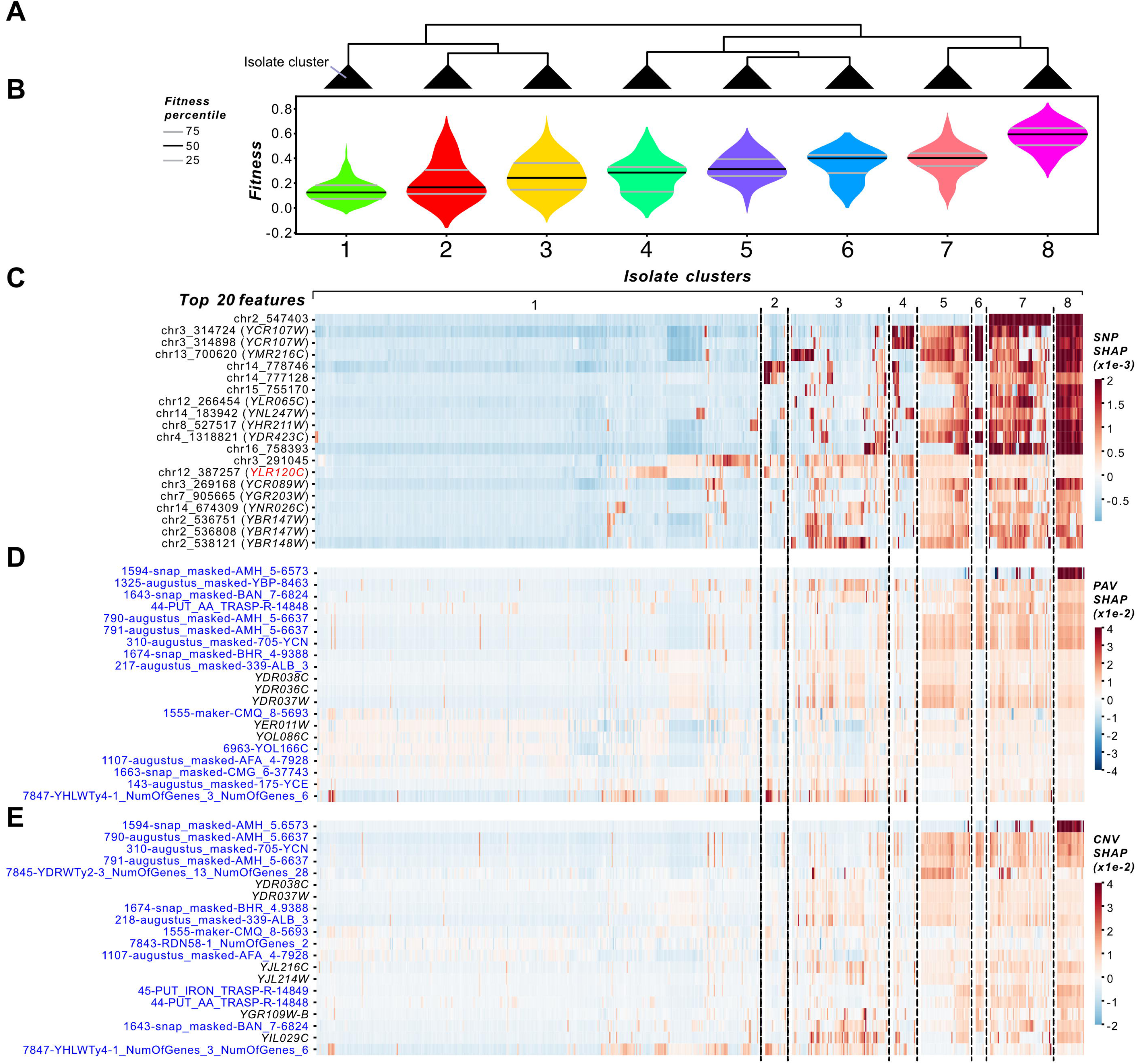
Isolate-dependent effects of SNP, PAV, and CNV features on fitness in YPD Caffeine 40 mM. **(A)** Dendrogram showing clusters of isolates based on the SHAP values of the top 20 features from the YPD Caffeine 40 mM optimized SNP model. **(B)** Violin plot of fitness distributions of isolates in each cluster identified in (A). Heatmap of SHAP values of the top 20 **(C)** SNP, **(D)** PAV, and **(E)** CNV features from the optimized YPD Caffeine 40 mM models. ORF features are colored in blue and features that are mapped to genes are colored in black on the left side of the heatmaps. Heatmap cells are colored by SHAP value. Isolates are ordered based on the SNP-based isolate clusters.

**S7 Fig.**
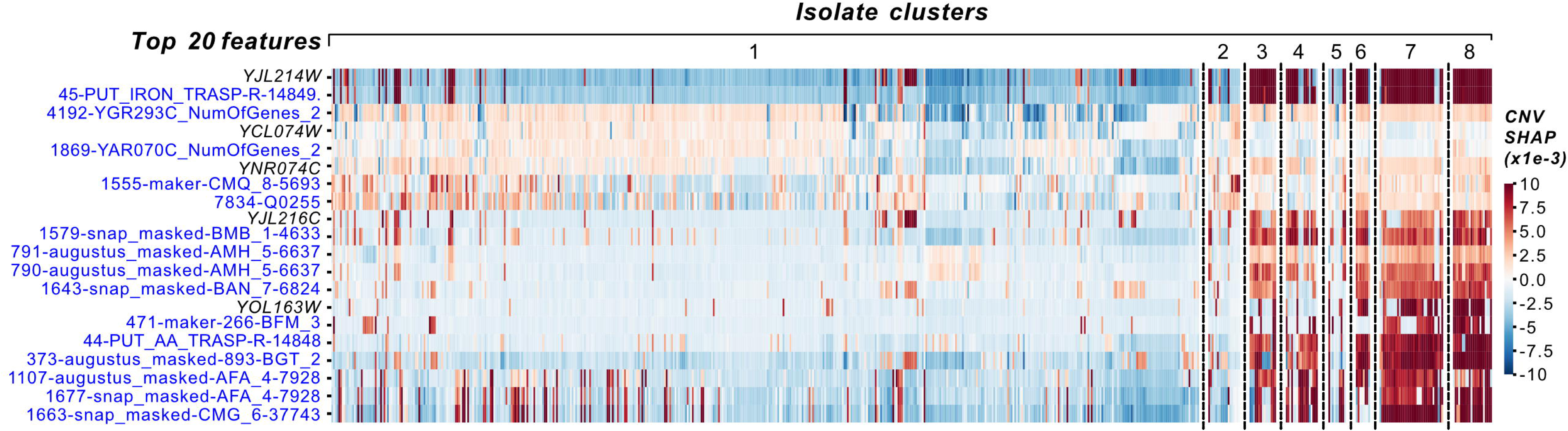
Isolate-based clusters of CNV feature SHAP values from the optimized YPD Benomyl 500 μg/ml RF model. Heatmap of SHAP values of the top 20 CNV features from the optimized YPD Benomyl 500 μg/ml RF model. ORF features are colored in blue and features that are mapped to genes are colored in black on the left side of the heatmap. Heatmap cells are colored by SHAP value and isolates are ordered based on the SNP-based isolate clusters.

**S8 Fig.**
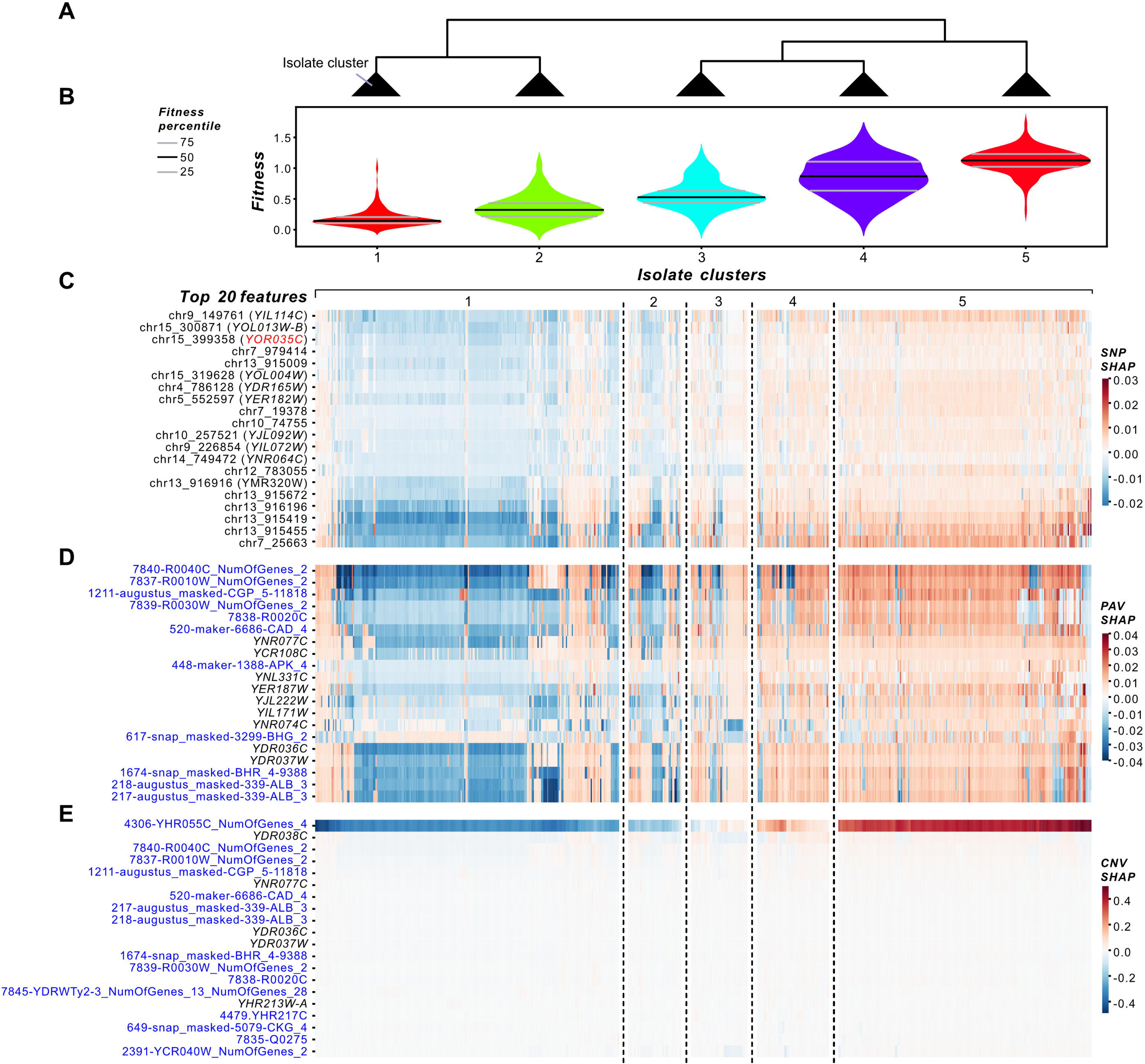
Isolate-dependent effects of SNP, PAV, and CNV features on fitness in YPD CuSO_4_ 10 mM. **(A)** Dendrogram showing clusters of isolates based on the SHAP values of the top 20 features from the YPD CuSO_4_ 10 mM optimized CNV model. **(B)** Violin plot of fitness distributions of isolates in each cluster identified in (A). Heatmap of SHAP values of the top 20 **(C)** SNP, **(D)** PAV, and **(E)** CNV features from the optimized YPD CuSO_4_ 10 mM models. ORF features are colored in blue and features that are mapped to genes are colored in black on the left side of the heatmaps. Heatmap cells are colored by SHAP value. Isolates are ordered based on the CNV-based isolate clusters.

**S9 Fig.**
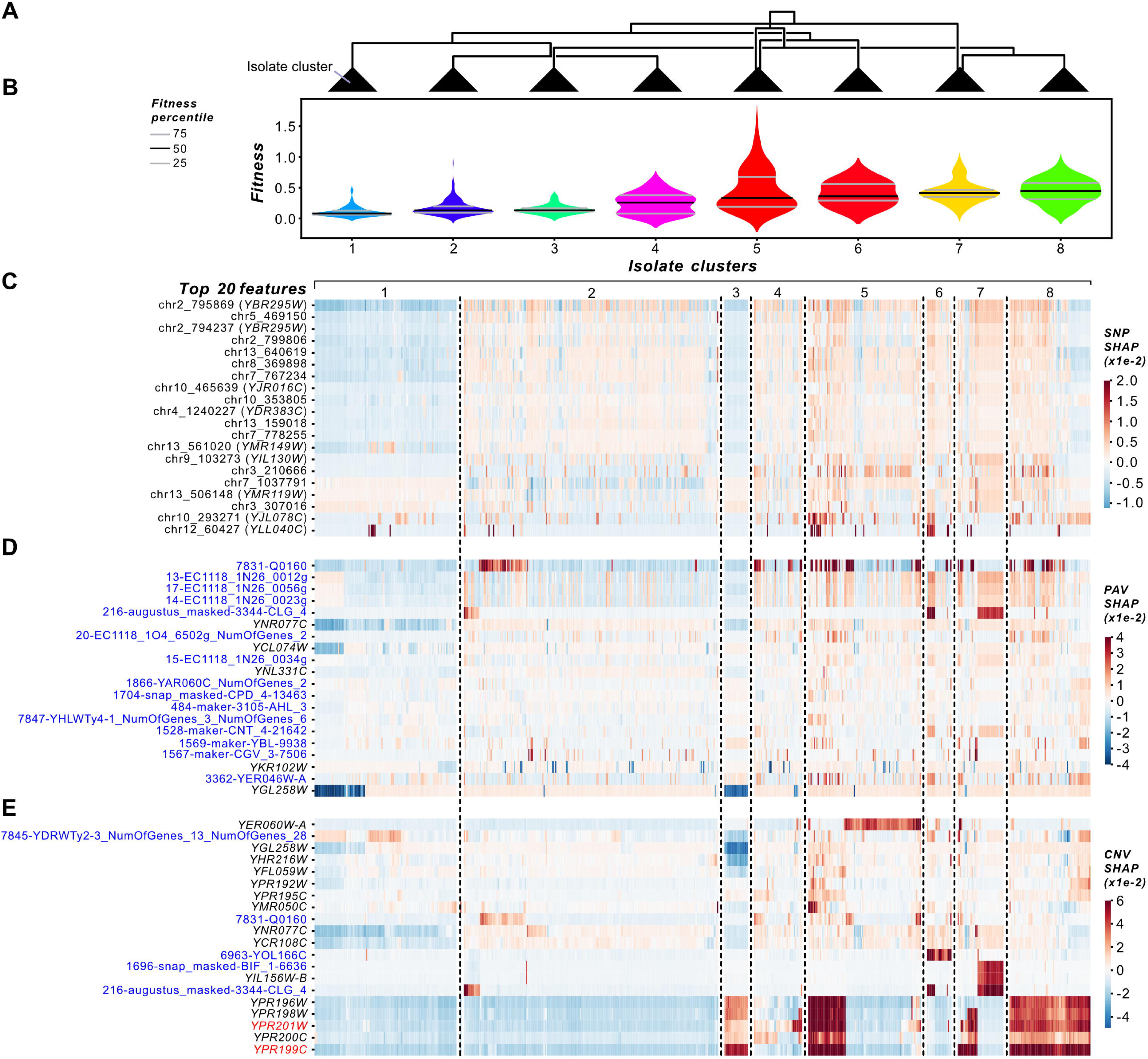
Isolate-dependent effects of SNP, PAV, and CNV features on fitness in YPD Sodium meta-arsenite 2.5 mM. **(A)** Dendrogram showing clusters of isolates based on the SHAP values of the top 20 features from the YPD Sodium meta-arsenite 2.5 mM optimized CNV model. **(B)** Violin plot of fitness distributions of isolates in each cluster identified in (A). Heatmap of SHAP values of the top 20 **(C)** SNP, **(D)** PAV, and **(E)** CNV features from the optimized YPD Sodium meta-arsenite 2.5 mM models. Benchmark gene systematic identifiers are colored in red, ORF features are colored in blue, and features that are mapped to genes are colored in black on the left side of the heatmaps. Heatmap cells are colored by SHAP value. Isolates are ordered based on the CNV-based isolate clusters.

**S10 Fig.**
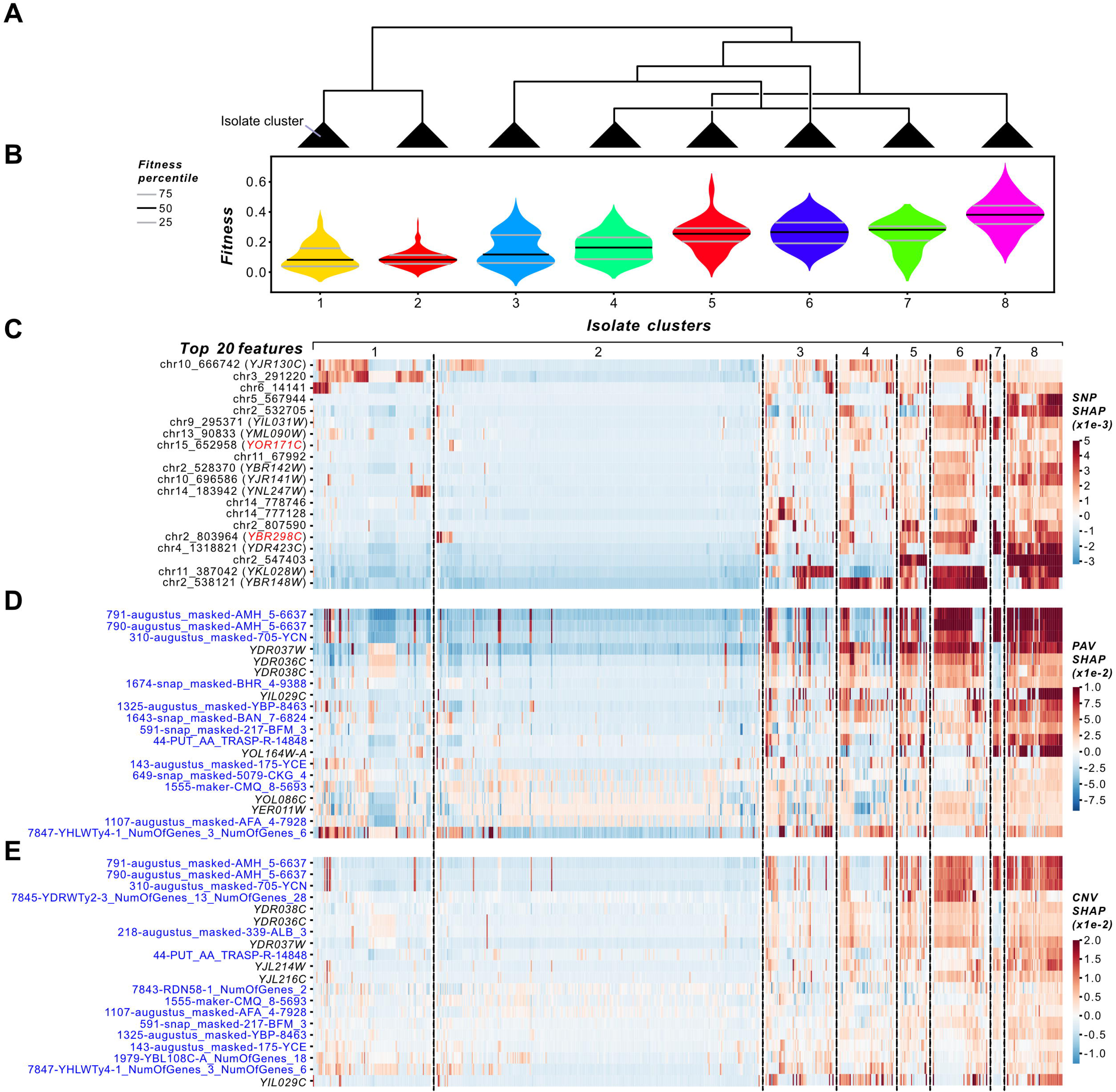
Isolate-dependent effects of SNP, PAV, and CNV features on fitness in YPD Caffeine 50 mM. **(A)** Dendrogram showing clusters of isolates based on the SHAP values of the top 20 features from the YPD Caffeine 50 mM optimized SNP model. **(B)** Violin plot of fitness distributions of isolates in each cluster identified in (A). Heatmap of SHAP values of the top 20 **(C)** SNP, **(D)** PAV, and **(E)** CNV features from the optimized YPD Caffeine 50 mM models. Benchmark gene systematic identifiers are colored in red, ORF features are colored in blue, and features that are mapped to genes are colored in black on the left side of the heatmaps. Heatmap cells are colored by SHAP value. Isolates are ordered based on the SNP-based isolate clusters.

**S11 Fig.**
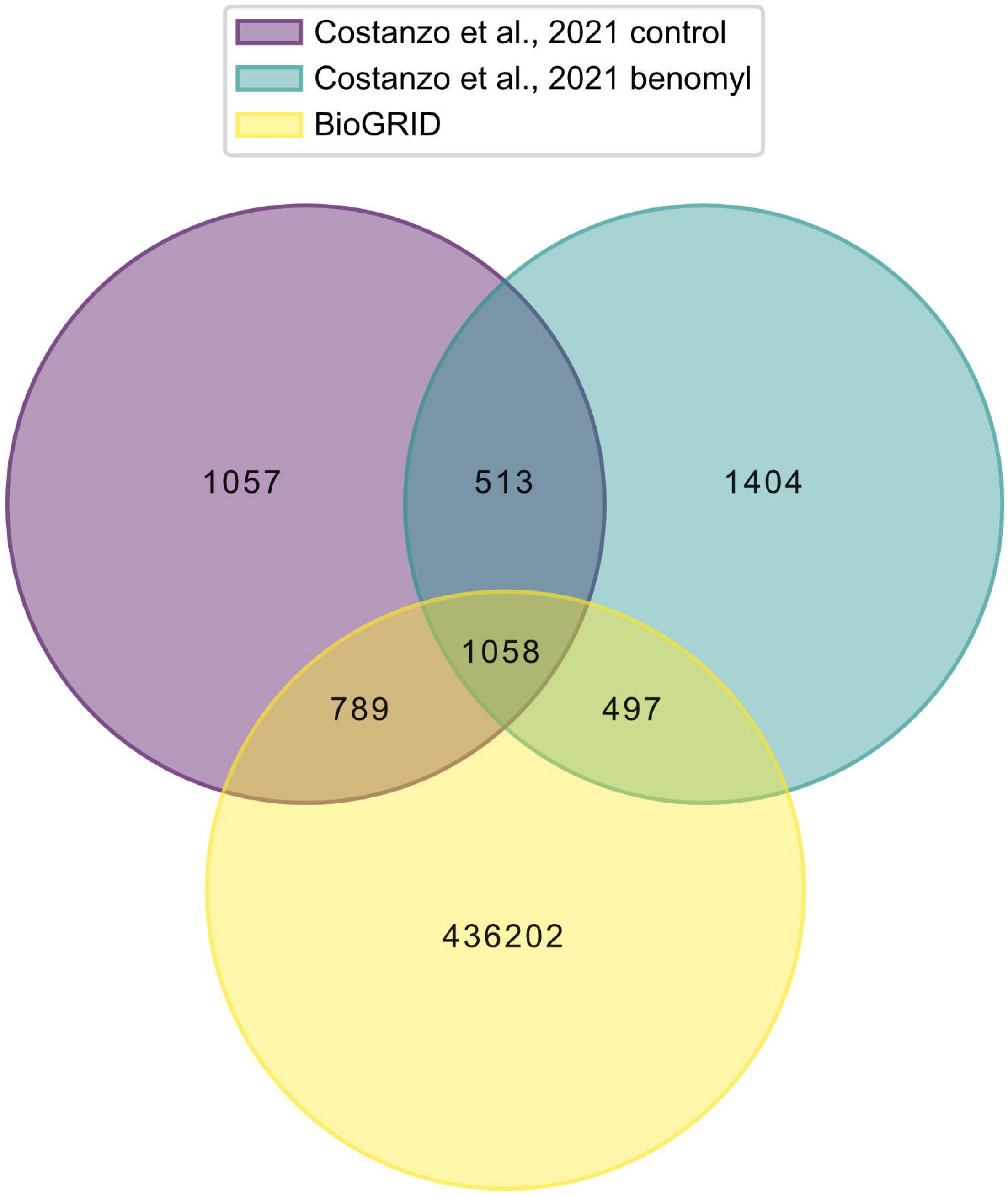
Experimentally validated genetic interactions from BioGRID and Costanzo et al., 2021. Venn diagram showing the overlap in genetic interactions collected from BioGRID (yellow) and genetic interactions validated by Costanzo et al., 2021 under the control condition (purple) and under 30 µg/ml benomyl (blue).

**S12 Fig.**
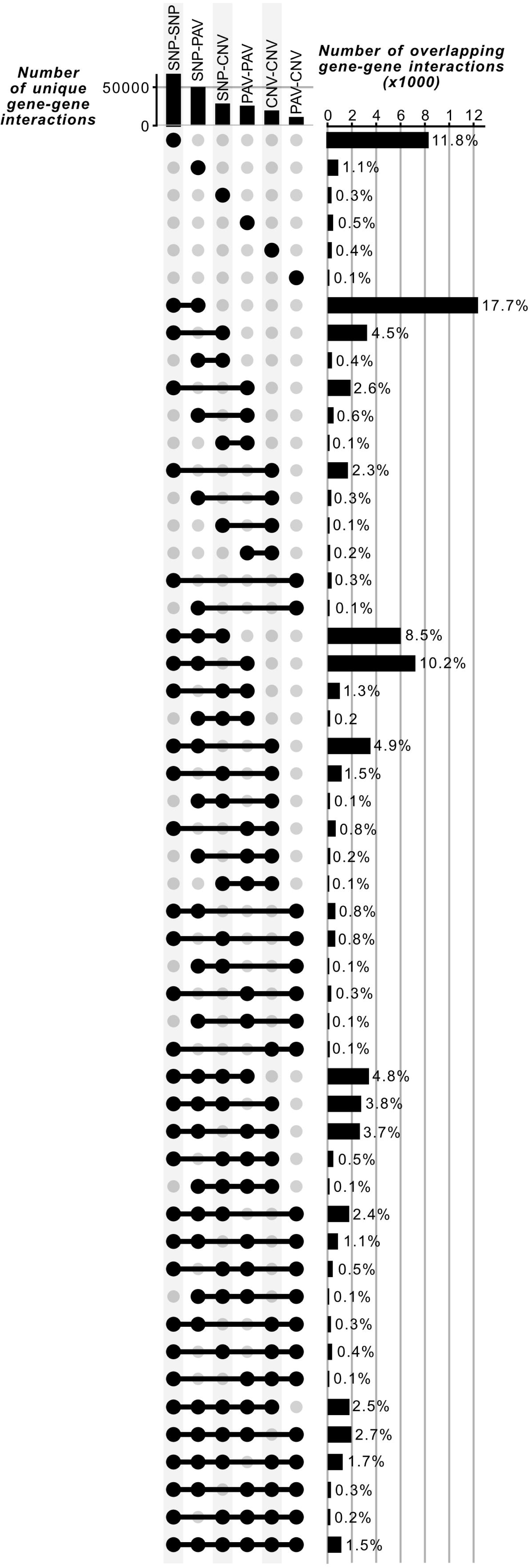
Comparison of unique gene-gene interactions represented by different variant-variant SHAP interactions. The number of unique (top-left bar chart) and overlapping gene-gene interactions (right bar chart) identified by SHAP interactions for different variant pair types. For the right bar chart, a single black dot represents the number of gene-gene interactions identified by a variant-variant interaction type (i.e., SNP-SNP, PAV-PAV, CNV-CNV, SNP-PAV, SNP-CNV, and PAV-CNV). The black line connecting two or more dots represents the comparison that is being made (e.g., comparing the overlap between SNP-SNP and PAV-PAV gene-gene interactions). The percentages of overlapping gene-gene interactions out of the total number of gene-gene interactions (69,486) are shown.

## Supplemental table legends

**S1 Table. List of environmental conditions.**

The list of 35 environments from which fitness values were obtained.

**S2 Table. Performances of single-environment models built using different algorithms and genetic variant features.**

For each of the 35 environments, a single-environment fitness prediction model was trained using linear and non-linear algorithms. Model training consisted of hyperparameter tuning via 5-fold cross-validation, training the model with the best hyperparameters within a 5-fold cross-validation scheme, and evaluating the model on a held-out test set consisting of one-sixth of the yeast isolates (125 out of 750 isolates). See **Materials and Methods** for details on model training. Models were trained on the first five principal components of the SNP features (PCs), SNPs, PAVs, or CNVs. Feature selection was implemented using the RF models and the subsequent optimized feature sets were used to train XGBoost, rrBLUP, Bayesian LASSO, and BayesC models. Reported metrics include average validation and test set performances, measured by R^2^ and Pearson’s correlation coefficient.

**S3 Table. Comparison of model performances by algorithm, environment, and genetic variant type.**

Pearson correlation coefficients of model performances across the 35 single-environment RF models trained using different algorithms. Correlations were calculated between pairs of algorithms, pairs of environments across algorithms or genetic variant types, and pairs of genetic variant types used to train models. A Mann-Whitney U test was also performed for pairs of algorithms to determine which algorithm generally performed the best.

**S4 Table. Contribution of fitness-related features and the number of features used to train models to RF model performances.**

Linear regression analysis of the model performances of the 35 PC, SNP, PAV, or CNV optimized RF models against three fitness-related features: narrow-sense heritability (*h^2^*), fitness variance, and median fitness in each environment, in addition to all pairwise and three-way interaction features (e.g., *h^2^*-by-fitness variance, see **Materials and Methods**). All features were centered and scaled prior to regression. SHAP values for each feature were estimated using LinearExplainer (see **Materials and Methods**). The feature values and SHAP values are provided in separate sheets. In summary, the variance in fitness in the five best predicted environments ranges from 0.014 to 0.210, and in the remaining environments, the variance ranges from 0.006 to 0.087. Median fitness values ranged from 0.11 to 0.51 for the five best predicted environments and 0.11 to 0.96 for the remaining environments. Fitness variance-by-*h^2^* interaction term values ranged from 0.011 to 0.192 for the five best predicted environments and 0.003 to 0.055 for the remaining environments. For SNPs, the term with the smallest *p*-value was *h^2^* (*P* = 0.06, coefficient = 0.07). An additional linear regression analysis was conducted using only the number of features used to train the optimized RF models.

**S5 Table. Comparison of feature ranks between Gini importance and SHAP values.**

Spearman’s rank correlation between Gini importance and average absolute SHAP values of features from single-environment RF models trained on SNPs, PAVs, or CNVs. Rho values, *p*-values, and the number of overlapping features identified by both feature importance measures are reported for all 35 environments (see **Materials and Methods**).

**S6 Table. Comparison of feature ranks between genetic variant types.**

Spearman’s rank correlation of Gini or SHAP-based feature importances between RF models trained on different genetic variant types (SNP vs PAV, SNP vs CNV, and PAV vs CNV). Comparisons were performed for both the RF models trained on either complete or optimized feature sets for all 35 environments. Rho values, *p*-values, and the number of overlapping genes identified by both feature importance measures are reported for all 35 environments (see **Materials and Methods**).

**S7 Table. Comparison of feature ranks between environments.**

Spearman’s rank correlation (rho) of rankings based on Gini or average absolute SHAP feature importances between environments. Correlations were calculated separately for SNP, PAV, and CNV single-environment RF models trained on either the optimized or complete feature sets. The number of overlapping features between environments are provided in a separate sheet.

**S8 Table. GO term and pathway enrichment analysis results.**

Enrichment of GO terms and pathway annotations (see **Materials and Methods**) among genes from the optimized single-environment RF models built using SNP, PAV, or CNV features. Enrichment analyses were performed separately for each genetic variant type, and genes from the optimized models were compared to a background set consisting of non-overlapping genes represented by the full set of SNP, PAV, or CNV features. Intergenic SNPs were excluded. Annotations were considered significantly enriched if they had a *q* < 0.05 after multiple testing correction using the Benjamini-Hochberg method to control the false discovery rate.

**S9 Table. SNP, PAV, and CNV feature importances.**

Gini importances or average absolute SHAP values of SNP, PAV, and CNV features from the optimized RF models for all 35 environments. Feature-to-S288C gene mappings are provided, along with GO term and pathway annotations and an indication of whether each gene is an SGD benchmark gene.

**S10 Table. Lists of experimentally validated benchmark fitness genes.**

Lists of benchmark genes experimentally proven to decrease fitness in the benomyl, caffeine, copper(II) sulfate, or sodium meta-arsenite environments. Benchmark genes were collected from SGD and manually curated from the literature.

**S11 Table. Enrichment analysis of benchmark genes.**

Enrichment analysis of benchmark genes from SGD for benomyl, caffeine, copper(II) sulfate, or sodium meta-arsenite stress and from the manually curated gene list. Results are reported for different rank percentile thresholds (ranks within the 1st, 5th, 10th, 15th, 20th, or 25th percentiles) for five environments: YPD Caffeine 40 mM, YPD Caffeine 50 mM, YPD Benomyl 500 μg/ml, YPD CuSO_ 10 mM, and YPD Sodium meta-arsenite 2.5 mM.

**S12 Table. The association between feature number and performance of benchmark gene models.**

A linear regression was performed using the number of features as the independent variable and the performance R^2^ of the test set as the dependent variable. Reported are the slope, standard error of the slope, intercept, *p*-value of the hypothesis test (alternative hypothesis: the slope of the regression line is nonzero), and the Pearson’s correlation coefficient.

**S13 Table. Effect of genetic relatedness to the genetic background used for experimental validation of benchmark genes on feature importance.**

SNP and PAV genotypes of the training isolates were compared to those of the laboratory strain S288C using Euclidean distance as a measure of genetic relatedness. K-means clustering was used to identify clusters of isolates. A Mann-Whitney U test was conducted to compare the median absolute SHAP values of benchmark genes between clusters of isolates. Results are reported for five environments: YPD Caffeine 40 mM, YPD Caffeine 50 mM, YPD Benomyl 500 μg/ml, YPD CuSO_ 10 mM, and YPD Sodium meta-arsenite 2.5 mM. See **Materials and Methods** for details on measuring genetic relatedness, cluster analysis, deriving gene-level feature importance, and conducting the Mann-Whitney U test.

**S14 Table. Correlation between fitness and SHAP values of isolates within a cluster for the top 20 predictive features.**

Isolates that were used to train optimized RF models were clustered based on the SHAP values of the top 20 features from the optimized SNP, PAV, or CNV RF models for five environments (YPD Caffeine 40 mM, YPD Caffeine 50 mM, YPD Benomyl 500 μg/ml, YPD CuSO_ 10 mM, and YPD Sodium meta-arsenite 2.5 mM). Within each cluster, the Pearson correlation coefficient between median absolute SHAP values, which was calculated across the top 20 features, and median fitness of isolates was quantified and a linear regression was fitted.

**S15 Table. ORF to S288C gene mappings based on BLAST.**

Table containing the systematic identifiers of S288C genes that were mapped to ORF features. For each ORF, the minimum E-value across BLASTx and tBLASTx alignments and the average percent identity are reported.

**S16 Table. Performance of models used to obtain SHAP interaction scores and its association with feature number.**

Performance of RF models built using reduced SNP, PAV, CNV, or integrated feature sets for predicting fitness in the YPD Benomyl 500_μg/mL environment (see **Estimating SHAP interaction scores**). A linear regression line was fitted to the model performances on the number of features used to train models. Linear regression results are provided in a separate sheet.

**S17 Table. Gene-gene interactions identified by the SHAP-based feature interactions.**

SHAP interaction scores from YPD Benomyl 500_μg/mL RF models trained using the reduced SNP, CNV, or integrated feature sets. SHAP interactions from the PAV model were excluded because of the poor performance on the test set.

**S18 Table. Experimentally validated gene-gene interactions identified by different variant-variant interaction types.**

The list of variant-variant feature interaction types are listed for each gene pair with experimental validation information.

**S19 Table. Enrichment of experimentally validated genetic interactions.**

Enrichment of experimentally validated genetic interactions in the SHAP interactions from the six RF models trained using the SNP, CNV, SNP + PAV, SNP + CNV, PAV + CNV, or SNP + PAV + CNV feature sets. Enrichment was assessed within the 1st, 5th, 10th, 15th, 20th, and 25th rank percentiles based on SHAP interaction scores for each model.

## Supplemental file legends

**S1 File. Fitness measurements.**

Fitness in 35 environments for 750 diploid *Saccharomyces cerevisiae* isolates.

**S2 File. SNP genotypes.**

Filtered SNP matrix containing 118,382 bi-allelic SNPs, which are encoded as -1, 0, or 1.

**S3 File. Kinship matrix.**

Kinship matrix from the SNP data.

**S4 File. PC matrix.**

First five principal components of the SNP data.The variance explained by each principal component is also provided.

**S5 File. PAV matrix.**

Filtered presence/absence variant data containing 7,708 ORFs.

**S6 File. CNV matrix.**

Filtered copy number variant data containing 7,708 ORFs.

**S7 File. Feature importances from the RF models built using complete feature sets.**

Gini importance and average absolute SHAP values of SNP, PAV, and CNV features from the single-environment RF models built using complete feature sets.

**S8 File. SNP-to-gene map.**

Table containing the systematic identifiers of S288C genes that were mapped to SNP features. Four additional columns indicate whether the S288C gene is a benchmark fitness gene for benomyl, caffeine, CuSO_4_, and/or sodium meta-arsenite.

**S9 File. ORF-to-gene map.**

Table containing the systematic identifiers of S288C genes that were mapped to ORFs. Four additional columns indicate whether the S288C gene is a benchmark fitness gene for benomyl, caffeine, CuSO_4_, and/or sodium meta-arsenite.

**S10 File. Experimentally validated genetic interactions.**

The combined set of 441,520 unique experimentally verified genetic interactions from the BioGRID database and the benomyl and control condition networks from Costanzo et al., 2016.

**S11 File. Performances of benchmark gene models.**

The number of features and validation/testing performance R^2^ values of RF models trained using benchmark gene sets, important non-benchmark gene sets, and combined benchmark + important non-benchmark gene sets for the YPD Caffeine 40_mM, YPD Caffeine 50_mM, YPD Benomyl 500_μg/mL, YPD CuSO_ 10_mM, and YPD Sodium meta-arsenite 2.5_mM environments (see **Assessing the contribution of benchmark genes to fitness predictions** for details).

**S12 File. SNP genotypes with S288C genotypes included.**

118,382 bi-allelic SNP genotypes encoded as 0 (homozygous for the reference allele), 1 (heterozygous), and 2 (homozygous for the alternative allele). S288C SNP genotypes were encoded as a vector of 0s.

**S13 File. Euclidean distance of the SNP genotypes.**

Genetic distances between the 625 training isolates and S288C, which were calculated using Euclidean distance of the SNP genotype matrix.

**S14 File. Euclidean distance of the PAV genotypes.**

Genetic distances between the 625 training isolates and S288C, which were calculated using Euclidean distance of the PAV genotype matrix.

**S15 File. SHAP cluster distance thresholds.**

Distance thresholds used to define isolate clusters at different granularities, based on the SHAP values of the top 20 SNP, PAV, and CNV features from the optimized YPD Caffeine 40 mM, YPD Caffeine 50 mM, YPD Benomyl 500_μg/mL, YPD CuSO_ 10 mM, and YPD Sodium meta-arsenite 2.5 mM RF models.

**S16 File. SHAP values of isolates clustered based on the top 20 important features.**

SHAP values of the top 20 SNP, PAV, and CNV features from the optimized RF models for YPD Caffeine 40 mM, YPD Caffeine 50 mM, YPD Benomyl 500_μg/mL, YPD CuSO_ 10 mM, and YPD Sodium meta-arsenite 2.5 mM. The row and column ordering of each SHAP value matrix corresponds to the heatmaps shown in Figs 5, S6, S7, S8, S9, and S10.

